# Human CST suppresses origin licensing and promotes AND-1/Ctf4 chromatin association

**DOI:** 10.1101/561977

**Authors:** Yilin Wang, Kathryn S. Brady, Benjamin Caiello, Stephanie M. Ackerson, Jason A. Stewart

**Affiliations:** Department of Biological Sciences, University of South Carolina, Columbia, SC, USA 29208; Center for Colon Cancer Research, University of South Carolina, Columbia, SC, USA 29208

**Keywords:** CDT1, DNA replication, MCM, origin licensing, replisome

## Abstract

Human CTC1-STN1-TEN1 (CST) is an RPA-like single-stranded DNA binding protein that interacts with DNA polymerase α-primase (pol α) and functions in telomere replication. Previous studies suggest that CST also promotes replication restart following fork stalling. However, the precise role of CST in genome-wide replication remains unclear. In this study, we sought to understand whether CST alters origin licensing and activation. Replication origins are licensed by loading of the minichromosome maintenance 2-7 (MCM) complex in G1 followed by replisome assembly and origin firing in S-phase. We find that CST directly interacts with the MCM complex and disrupts binding of CDT1 to MCM, leading to decreased origin licensing. We also show that CST enhances replisome assembly by promoting AND-1/pol α chromatin association. Moreover, these interactions are not dependent on exogenous replication stress, suggesting that CST acts as a specialized replication factor during normal replication. Overall, our findings implicate CST as a novel regulator of origin licensing and replisome assembly/fork progression through interactions with MCM, AND-1 and pol α.

## Introduction

DNA replication must occur with high fidelity and efficiency to preserve genome stability. Each time human cells divide 50,000-100,000 DNA replication origins are activated for genome duplication (Higa, Fujita et al., 2017, Riera, Barbon et al., 2017, Zhai, Li et al., 2017b). During telophase and G1, replication origins are licensed by binding of the origin recognition complex (ORC) and CDC6 to the DNA, followed by recruitment of CDT1 and minichromosome maintenance 2-7 (MCM) to form the pre-replication complex (pre-RC). Loading of the first MCM hexamer by ORC and CDC6 leads to the ORC-CDC6-CDT1-MCM (OCCM) complex. A second MCM hexamer is then recruited and loaded onto the DNA for origin licensing. Recruitment and loading of MCMs are dependent on CDT1. CDT1 facilitates interaction between MCM and ORC-CDC6 and also stabilizes opening of the MCM hexamer for loading onto the DNA (Frigola, He et al., 2017, Masai, Matsumoto et al., 2010, Pozo & Cook, 2016). Once the first MCM hexamer is loaded, CDT1 and CDC6 are released. A second MCM-CDT1 complex along with CDC6 then bind ORC, leading to loading of a second MCM hexamer (Ticau, Friedman et al., 2017, Ticau, Friedman et al., 2015). Loading of the two MCM hexamers constitutes a licensed replication origin. Origin licensing is restricted to telophase and G1 of the cell cycle to prevent re-replication in S-phase. Unlike budding yeast, origin licensing in mammals is not defined by DNA sequence but by chromatin context and accessibility (Cayrou, Ballester et al., 2015).

Upon entering S-phase, replication factors are recruited to origins to form the pre-initiation complex (pre-IC). MCM is bound by CDC45 and GINS to form the CDC45-MCM-GINS (CMG) complex, which serves as the replicative helicase (Deegan & Diffley, 2016). Three DNA polymerases (pol) are recruited during replisome assembly and used for DNA synthesis upon origin firing. Pol ε binds directly to CMG, while pol δ and pol α-primase (pol α) are linked to the replisome by PCNA and Ctf4/AND-1, respectively. Once assembled, the replisome is then activated, or fired, following phosphorylation of MCM by Dbf4-dependent kinase (DDK) and cyclin-dependent kinase (CDK). Origin licensing and activation was recently reconstituted with purified replication factors from budding yeast (Yeeles, Deegan et al., 2015). However, many questions remain, particularly in regards to where replication origins are licensed in higher eukaryotes and how they are selected for activation. Here, we identify human CTC1-STN1-TEN1 (CST) as a novel regulator of origin licensing and replisome assembly. CST is an RPA-like ssDNA binding protein that has primarily been characterized as a telomere replication factor with less well understood roles in genome-wide replication (Stewart, Wang et al., 2018).

Our previous work indicated that CST promotes origin firing in response to genome-wide replication stress (Stewart, Wang et al., 2012). In addition, work by Chastain et al. showed that CST recruits RAD51 to rescue stalled replication and prevent chromosome fragility at GC-rich DNA (Chastain, Zhou et al., 2016). However, the mechanism by which CST facilitates replication restart remains unclear. CTC1 and STN1 were originally discovered as pol α accessory factors (Casteel, Zhuang et al., 2009, Goulian, Heard et al., 1990). CST stimulates pol α primase activity and the primase-to-polymerase switch (Ganduri & Lue, 2017, Nakaoka, Nishiyama et al., 2012). Nevertheless, CST does not localize to active replication forks, suggesting it may function prior to replication initiation and/or at stalled replication forks (Miyake, Nakamura et al., 2009, Sirbu, McDonald et al., 2013). Moreover, depletion of CST subunits does not alter bulk DNA replication under normal conditions but does result in increased anaphase bridges and chromosome fragility, suggesting that CST is likely utilized at specific regions of the genome (Chastain et al., 2016, Stewart et al., 2012, Wang, Stewart et al., 2012, Wang & Chai, 2018).

In agreement with this idea, *in vitro* biochemical analysis revealed that CST binds and resolves G-quadruplexes (G4s) (Bhattacharjee, Wang et al., 2017). ChIP-seq analysis also demonstrated that STN1 localizes to non-telomeric GC-rich regions, which are known to form G4s (Chastain et al., 2016). G4s are stable, four-stranded structures that can block replication, regulate transcription and are associated with several diseases (Maizels, 2015, Rhodes & Lipps, 2015). G4s are also enriched at DNA replication origins and may promote origin licensing (Valton & Prioleau, 2016).

During telomere replication, CST participates in many of the steps required for telomere maintenance. These steps include replication of the telomere duplex, removal of telomerase, prevention of ATR activation and engagement of pol α for C-strand fill-in synthesis (Chen, Redon et al., 2012, Feng, Hsu et al., 2018, Feng, Hsu et al., 2017, Gu & Chang, 2013, Gu, Min et al., 2012, Kasbek, Wang et al., 2013, Miyake et al., 2009, Stewart et al., 2012, Surovtseva, Churikov et al., 2009, Wang et al., 2012). CST was also recently shown to interact with the shieldin complex to counteract double-strand break (DSB) end resection by facilitating fill-in by pol α, similar to its role in telomeric C-strand fill-in (Barazas, Annunziato et al., 2018, Mirman, Lottersberger et al., 2018). This multi-functionality of CST in telomere maintenance, DSB repair and replication rescue is not unexpected given its similarity to RPA.

RPA directs various transactions during DNA replication and repair through the use of multiple oligonucleotide-oligosaccharide binding (OB)-folds, which allow RPA to bind ssDNA of different lengths and configurations (e.g. ss-dsDNA junctions) (Chen & Wold, 2014, Fanning, Klimovich et al., 2006). Like RPA, CST is composed of multiple OB-folds, which allow it to bind different form of ssDNA (Bhattacharjee et al., 2017, Chen et al., 2012, Miyake et al., 2009). However, while CST and RPA have similar binding affinities, CST, unlike RPA, is in low abundance and may require recruitment to specific sites. For example, recruitment of CST to DSBs is dependent on the shieldin complex, whereas TPP1 appears to localize CST to telomeres (Chen et al., 2012, Mirman et al., 2018). Use of CST may be advantageous because, unlike RPA, its binding does not elicit ATR activation (Feng et al., 2017).

Mutations in CTC1 and STN1 cause two pleiotropic autosomal recessive disorders, Coats plus and dyskeratosis congenita (Anderson, Kasher et al., 2012, Keller, Gagne et al., 2012, Polvi, Linnankivi et al., 2012, Simon, Lev et al., 2016). Dyskeratosis congenita is caused by accelerated telomere shortening, which leads to cell proliferation defects and ultimately bone marrow failure (Armanios & Blackburn, 2012, Stanley & Armanios, 2015). Coats plus shares features of dyskeratosis congenita but has additional features, including intercranial calcifications, retinopathy, intrauterine growth retardation and gastrointestinal bleeding (Briggs, Abdel-Salam et al., 2008). Interestingly, some Coats plus patients do not exhibit accelerated telomere shortening, suggesting that both telomere and non-telomere defects contribute to the disease (Kotsantis, Petermann et al., 2018, Polvi et al., 2012, Romaniello, Arrigoni et al., 2013). Thus, determining the different roles of CST in preserving genome stability is necessary to characterize the molecular etiology of these diseases.

Here, we present evidence that CST interacts with the MCM complex and suppresses origin licensing by disrupting the interaction between MCM and CDT1. Furthermore, we find that CST is important for recruitment/stabilization of AND-1 and pol α on the chromatin. Interestingly, regulation of both origin licensing and AND-1/pol α chromatin association occur in the absence of HU-induced replication fork stalling, suggesting that these functions are independent of CST’s role in origin activation following exogenous replication stress. Together, our results support a direct role for CST in both origin licensing and replication activation.

## Results

### STN1depletion leads to increased levels of chromatin-bound MCM

Since the precise role(s) of CST in general replication are poorly understood, we tested whether CST affects the levels of chromatin-bound MCM in the absence of replication stress, as this would suggest changes in origin licensing and activation. To determine whether depletion of STN1 alters MCM levels, we used previously characterized HeLa1.2.11 cells with stable shRNA STN1 knockdown (shSTN1) to measure the levels of chromatin-bound MCM (Figure 1) (Stewart et al., 2012). Cells expressing a non-targeting shRNA (shNT) or shSTN1 cells expressing an shRNA resistant Flag-STN1 (shSTN1-Res) were used as controls (Figure 1A). EdU was added to the cultures 30 min prior to sample processing, which enabled later identification of S-phase cells (see Figure 2A). Cells were then harvested, pre-extracted to remove soluble MCM, fixed and examined by indirect immunofluorescence (IF) to detect chromatin-bound MCM7, MCM3 or MCM6 (Figure 1B-C). MCM fluorescence intensity was measured in each nucleus and the MCM signal intensity compared between cell lines (Figure 1C). The analysis revealed that the levels of chromatin-bound MCM7, MCM3 and MCM6 were significantly increased in shSTN1 cells compared to controls. These results were confirmed in HCT116 shSTN1 cells for chromatin-bound MCM6 (Figure EV1A-B). The increase is specific to chromatin-bound MCM, as total cellular MCM3, MCM6 or MCM7 levels were similar in shSTN1 and controls cells (Figure EV1C). These findings indicate that depletion of STN1 significantly increases chromatin-bound MCM.

**Figure 1.**
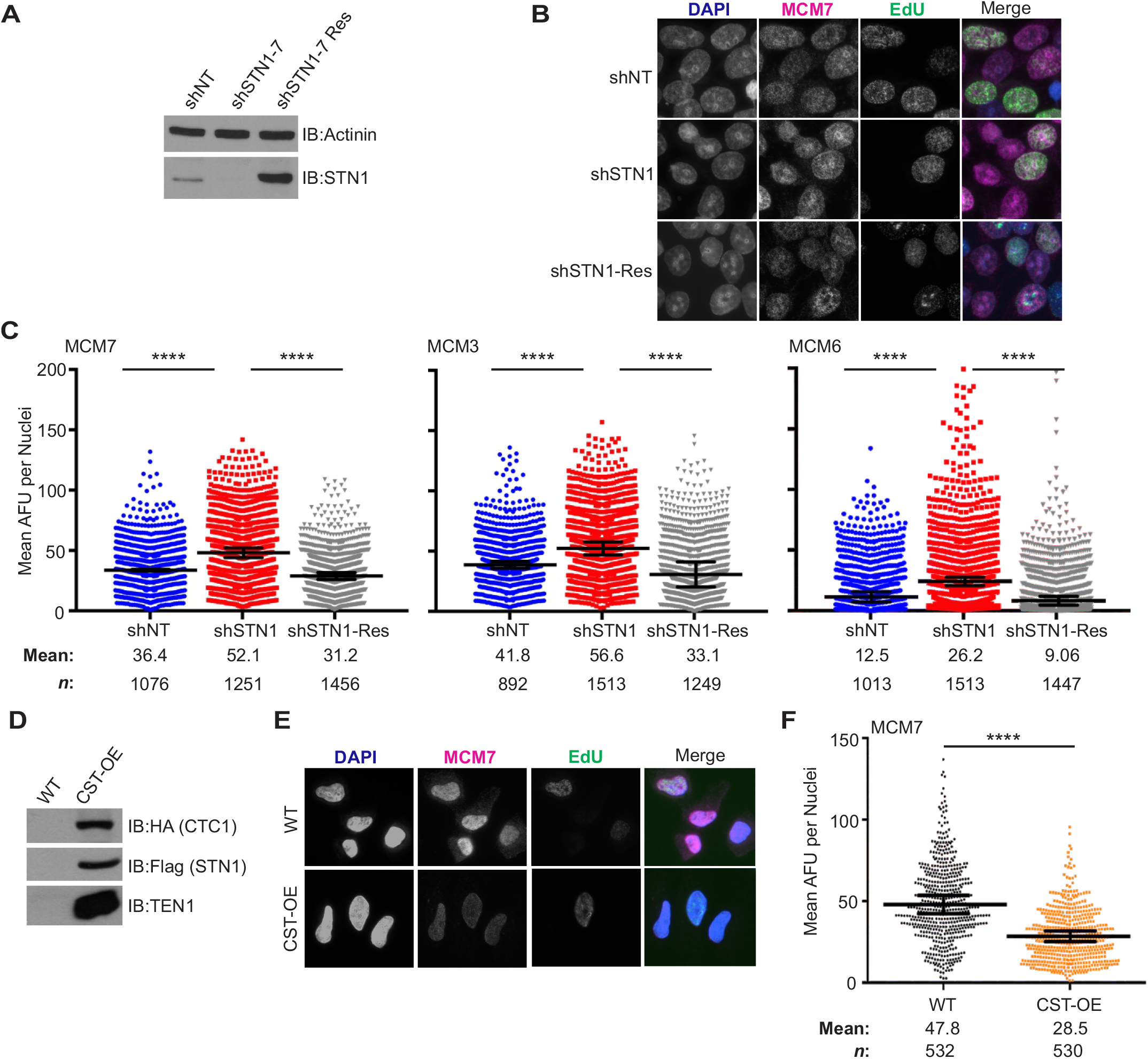
Altered CST expression leads to changes in chromatin-bound MCM. (**A**) Western blot analysis of STN1 knockdown in HeLa cells. Actinin is used as a loading control. (**B**) Representative images of pre-extracted, EdU-lableled cells used to measure MCM levels. shNT: non-targeting shRNA; shSTN1: STN1 shRNA; shSTN1-Res: shSTN1 cells plus shRNA-resistant Flag-STN1. DAPI: blue, MCM: magenta, EdU: green. (**C**) Dot plots of mean MCM7, MCM3 or MCM6 intensity per nuclei, represented in arbitrary fluorescent units (AFU). Black line and numbers below the graph indicate the mean AFU. Error bars indicate the +/-SEM of at least three independent biological experiments. *n*=indicates the number of total nuclei scored. (**D**) Western blot analysis of HA-CTC1, Flag-STN1 and untagged TEN1 in CST overexpressing (CST-OE) and wild type (WT) cells. (**E**) Representative images of HeLa WT and CST-OE cells, as in B. (**F**) Dot plots of mean MCM7 intensity per nuclei. Black line and numbers below the graph indicate the mean AFU. Error bars indicate the +/-SEM of three independent biological experiments. *n*=indicates the number of total nuclei scored. *P*-values were calculated by an unpaired, two-tailed Mann-Whitney test (\*\*\*\**p≤*0.0001).

**Figure 2.**
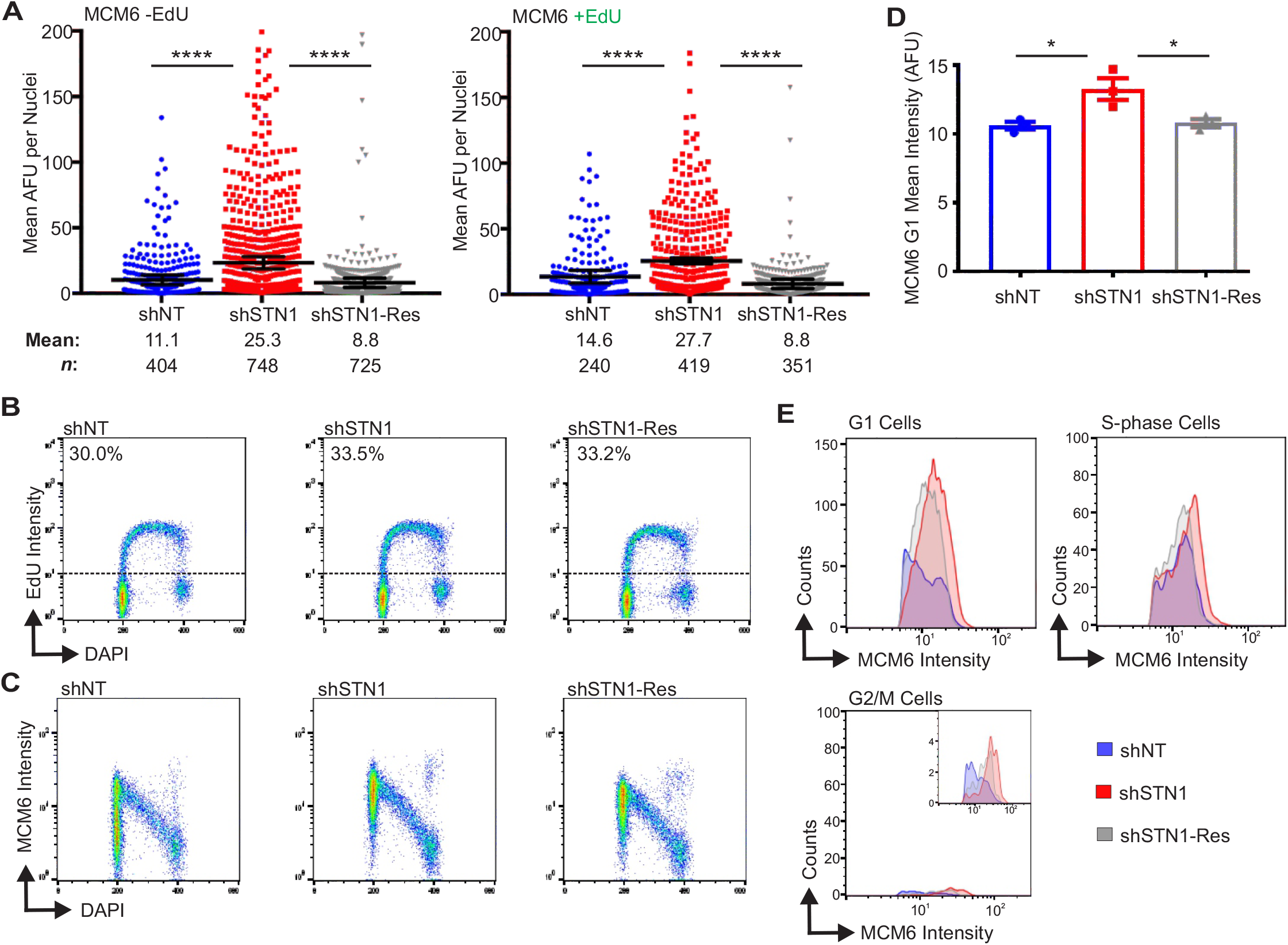
Chromatin-bound MCM increases in both G1 and S-phase with STN1 depletion. The indicated cell lines were labeled with EdU, pre-extracted and MCM6 detected. (**A**) Dot plots of mean MCM6 intensity. S-phase cells are indicated as +EdU. Black line and numbers below the graph indicate the mean AFU. Error bars indicate the +/-SEM of three independent biological experiments. *n*=indicates the number of total nuclei scored. (**B-E**) Flow cytometry data is representative of four independent biological experiments. (**B**) DNA content (DAPI) versus EdU signal intensity. Numbers above the dashed line represents the percentage of EdU+ cells. (**C**) DNA content versus MCM6 intensity. (**D**) Graph of the intensity of G1 MCM6 positive cells (see Materials and Methods and Figure EV2 for gating). (**E**) Histograms showing the distribution of MCM signal intensity for G1, S-phase or G2/M cells (see Materials and Methods). *P*-values were calculated by an unpaired, two-tailed Mann-Whitney test in (A) and student’s t-test in (D) (\*\*\*\**p≤*0.0001, * *p≤*0.05).

### CST overexpression leads to decreased chromatin-bound MCM

We next tested whether CST overexpression (CST-OE) also affected MCM chromatin association. For this experiment, we used a previously described HeLa cell line overexpressing all three CST subunits (Figure 1D) (Wang, Stewart et al., 2014). IF was performed, as described above, for either MCM7 (Figure 1E-F) or MCM6 (Figure EV1D). CST-OE had the reverse effect to STN1 depletion as it led to a substantial decrease in chromatin-bound MCM. Together, the results in Figure 1 indicate that the level of chromatin-bound MCM is inversely proportional to CST expression. It is notable that the changes in MCM occur in the absence of exogenous replication stress, which indicates that CST is needed to regulate MCM under normal conditions rather than in response to genome-wide replication fork stalling or DNA damage.

### STN1 depletion leads to increased MCM in G1 and S-phase

We next wanted to determine whether the changes in chromatin-bound MCM were due to alterations in loading of MCM in G1 or unloading in S/G2. MCM loading onto the chromatin is tightly regulated throughout the cell cycle with origin licensing restricted to telophase and G1. MCM is then unloaded from the chromatin following DNA synthesis in a ubiquitin-dependent manner (Dewar, Budzowska et al., 2015, Dewar, Low et al., 2017, Moreno, Bailey et al., 2014, Sonneville, Moreno et al., 2017). In initial experiments, we determined the effect of STN1 knockdown on the levels of chromatin-bound MCM in EdU positive versus negative cells (Figure 1B and 2A). This allowed us to distinguish whether the increase was restricted to S-phase. We found that MCM6 levels significantly increased in both EdU positive and EdU negative shSTN1 cells, indicating that the increase in MCM occurs both outside and within S-phase (Figure 2A).

Since the above experiments did not distinguish between cells in G1 versus G2, we next performed flow cytometry to separate MCM positive cells in G1 and G2/M populations. This allowed us to address whether the changes in chromatin-bound MCM reflected increased MCM loading (i.e. origin licensing) in G1 versus MCM dissociation in S/G2. HeLa control and shSTN1 cells were labeled with EdU for 30 min, pre-extracted to remove soluble MCM, stained for MCM6 and EdU and then separated by flow cytometry for cell cycle analysis (Figure EV2A). Cells were then gated to separate G1, S and G2/M populations (Figure EV2B-D and see Materials and Methods). The analysis revealed no observable change in the number or intensity of EdU positive cells with STN1 depletion compared to controls, suggesting DNA synthesis is not significantly affected with STN1 depletion, as previously reported (Figure 2B) (Wang et al., 2014, Wang et al., 2012). Likewise, when we compared chromatin-bound MCM across the cell cycle, we found the expected increase in MCM positive cells in G1 (origin licensing) followed by a linear decrease throughout S-phase (origin firing/DNA synthesis) with very few MCM6 positive cells in G2/M (Figure 2B) (Matson, Dumitru et al., 2017). However, the G1 population of shSTN1 cells exhibited an increase in MCM6 intensity compared to the control cells in mean intensity of MCM6 positive cells, suggesting increased origin licensing after STN1 depletion (Figure 2C-E). There was also an increase in MCM intensity in S-phase and G2/M (Figure 2E). However, very few MCM positive cells were within the G2/M population, which was unchanged with STN1 depletion (Figure 2E, MCM positive cells in G2/M: shNT=1.5%, shSTN1=1.5%, shSTN1-Res=1.4%). These data argue against a defect in MCM removal following replication termination because this should lead to an increase in the fraction of MCM positive cells in G2/M. Instead, our results demonstrate that CST suppresses origin licensing in G1. Similar results were observed in HCT116 shSTN1 cells (Figure EV2F-H). Overall, these results provide evidence that CST regulates origin licensing (i.e. MCM loading in G1). Furthermore, we find that chromatin-bound MCM increases in S-phase, which is likely caused, at least in part, by excess origin licensing in G1.

### CST interacts with the MCM complex

Since CST affects origin licensing, we next tested whether CST interacts with MCM (Figure 3). First, co-immunoprecipitation (co-IP) experiments were performed using whole cell lysates from cells expressing CST subunits. HA-tagged CTC1, Flag-tagged STN1, or untagged TEN1 were transiently expressed in combination or individually. The lysates were nuclease-treated prior to IP to ensure that the interaction was not DNA-dependent. Association of MCM subunits or CDC45 with CST was then assessed by Western blot (Figure 3A). MCM4 and MCM7 co-immunoprecipitated with CTC1 or STN1 (Figure 3A). CDC45, a component of the CMG complex, was also detected. Interaction with MCM was confirmed by reciprocal co-IP, where endogenous MCM7 was pulled down and STN1 detected (Figure 3B). Co-IP was also performed following HU treatment to see if interactions between CST and MCM or CDC45 was increased upon HU-induced fork stalling. However, neither interaction was significantly altered in response to HU treatment (Figure EV3A-B).

**Figure 3.**
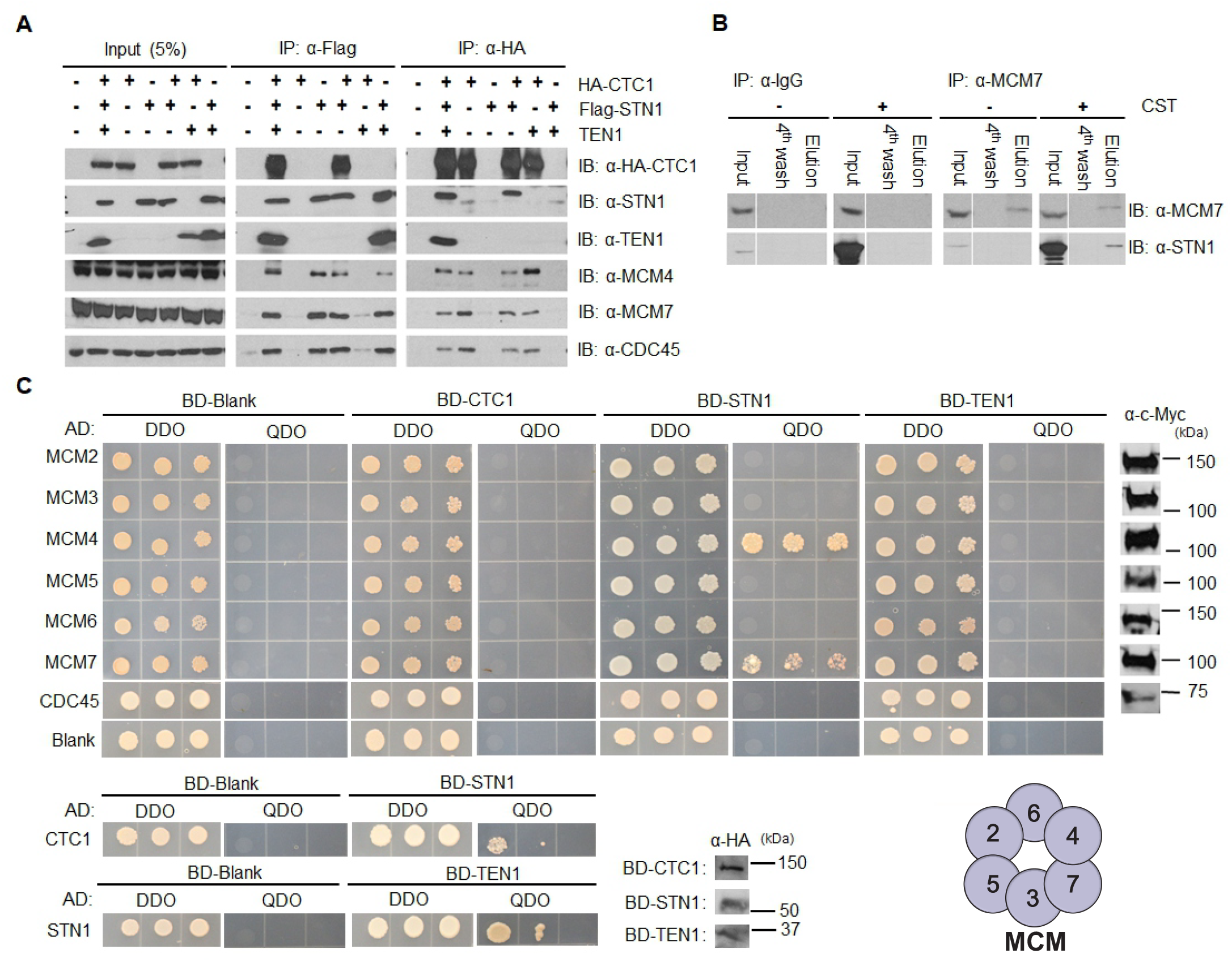
CST and MCM physically interact. (**A**) Co-immunoprecipitation (IP) of lysates from either mock or transfected HEK 293T cells with either Flag or HA antibody, as indicated. Data is representative of three independent biological experiments. (**B**) Reciprocal co-IP experiment with MCM7 or an IgG control antibody of lysates from HEK 293T cells transfected with plasmids expressing CST. Data is representative of three independent biological experiments. (**C**) Yeast-two-hybrid analysis of the interaction between CST and MCM. Double synthetic dropout (DDO) medium was used for plasmid selection and quadruple dropout (QDO) medium to assess interaction. Colony growth was recorded 3-4 days after plating. Interactions between CTC1 and STN1, STN1 and TEN1 were used as a positive control (bottom). Data is representative of three independent biological experiments. Expression of MCM subunits and CDC45 were detected by Western blot using a Myc antibody (far right). Expression of CST subunits was detected by Western blot using HA antibody (bottom middle). Representation of MCM hexamer with numbers to indicate subunits (bottom right).

Next, yeast-two-hybrid analysis was performed to more closely examine the interaction between CST and MCM, and to identify which subunit(s) interact. Human CTC1, STN1 or TEN1 coding regions were fused with the *GAL4* DNA-binding domain (BD), while human MCM subunits or CDC45 were linked to the *GAL4* DNA-activation domain (AD). Combinations of the constructs were then transformed into yeast, selected and spotted onto double synthetic dropout (DDO) medium, lacking histidine and tryptophan, or quadruple dropout (QDO) medium, lacking tryptophan, leucine, adenine and histidine. The DDO media was used to select for plasmid transformations and QDO media for cells producing adenine and histidine, which indicates protein interaction. Western blot of whole cell lysate was also performed to confirm the expression of MCM subunits, CDC45 and CST subunits (Figure 3C).

Among the different MCM subunits, we identified strong interactions between STN1 with MCM4 and MCM7 (Figure 3C). A weak interaction was also observed between CTC1 and MCM4 on the less stringent triple synthetic dropout (TDO) medium (Figure EV3C). TEN1 did not interact with any MCM subunit. Together, our results identify a direct interaction between STN1 and the MCM4-MCM7 interface. Interestingly, this region sits opposite the MCM2-MCM5 gate, suggesting that CST would not block opening of the gate or binding of CDC45 and GINS (Riera et al., 2017, Zhai et al., 2017b). We also tested CDC45 and CST subunits for interaction in the yeast-two-hybrid assay and did not observe any growth on the TDO or QDO media, which suggest that their association, detected by co-IP, may be bridged by MCM (Figure 3C). Alternatively, multiple CST subunits could be required for interaction, as is the case for CST interaction with pol α (Bhattacharjee et al., 2017, Feng et al., 2018). Overall our results demonstrate that CST interacts with MCM, with strong interaction between STN1 and MCM4-MCM7. Based on the yeast-two-hybrid data, we propose that this interaction is direct. However, it is possible that an evolutionary-conserved protein could bridge the interaction.

### CST disrupts the interaction between MCM and CDT1

We next sought to determine whether CST binding to MCM was the underlying cause of decreased origin licensing by CST. As mentioned previously, CDT1 plays an essential role in the recruitment and loading of MCM. Since CST interacts with MCM, we hypothesized that binding of CST could obstruct or destabilize CDT1 binding to MCM. Recent biochemical analysis of budding yeast MCM and Cdt1 showed that Cdt1 interacts with MCM2 and MCM6, and weakly with MCM4 (Frigola et al., 2017). Human CDT1 has also been shown to interact with MCM6 (Liu, Wu et al., 2012). To more closely examine the interaction between MCM and CDT1 in humans, we performed yeast-two-hybrid analysis (Figure 4A). In agreement with previous studies, we observed a strong interaction between CDT1 and MCM6. Additionally, we find a weaker interaction between CDT1 and MCM4, similar to budding yeast Cdt1. These findings suggest that CST and CDT1 have overlapping binding surfaces (i.e. CST binds MCM4-MCM7 while CDT1 binds MCM6-MCM4).

**Figure 4.**
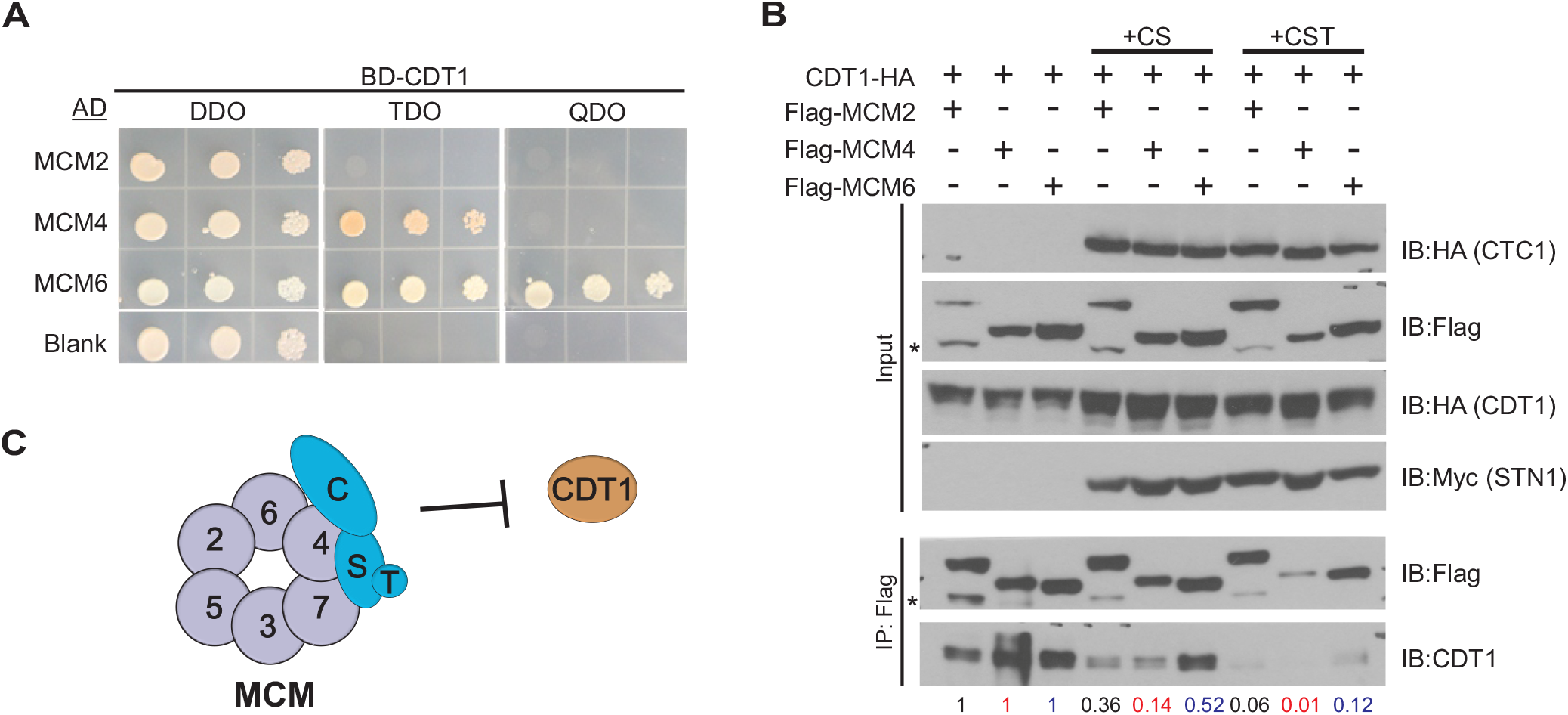
CST disrupts the interaction between MCM and CDT1. (**A**) Yeast-two-hybrid analysis was used to analyze the interaction between CDT1 and MCM. Yeast diploids growing on double synthetic dropout (DDO) medium were selected on triple dropout medium (TDO) or quadruple dropout (QDO) medium for binding and expression. Colony growth was monitored 3-4 days after plating and incubation. Data is representative of two independent biological experiments. (**B**) Co-IP of lysates from HEK 293T cells expressing Flag-MCM subunits and CDT1-HA with or without expression of CST subunits (+CS: CTC1 & STN1; +CST: CTC1, STN1, TEN1), as indicated. Relative CDT1-HA band intensity was then compared between no CST or CST expression, as indicated below the gel (see Material and Methods). Data is representative of three independent biological experiments for +CST. Replicate experiment is shown in Figure EV4. (**C**) Model of how CST binding to MCM could block CDT1 binding to MCM.

To test whether CST disrupts CDT1 binding to MCM, CDT1-HA, Flag-MCM subunits and/or CST were expressed in HEK 293T cells. Flag-tagged MCM2, MCM4 or MCM6 was then immunoprecipitated. The levels of CDT1-HA associated with MCM were then measured with or without expression of the entire CST complex (+CST) or just CTC1 and STN1 (+CS) (Figure 4B and EV4). As expected, CDT1 associated with MCM4 and MCM6 in the control (no CST or CS). Association was also observed with MCM2. However, when CST was co-expressed, we observed a decrease in CDT1 levels, suggesting that CST disrupts MCM-CDT1 interaction.

One caveat was that expression of MCM subunits and CDT1 varied to some extent across samples. To account for these differences, we normalize to the amount of MCM pulled down in +CST or +CS samples to MCM IPs without CST expression (see Materials and Methods for more details). Furthermore, we normalized the levels of CDT1 in the MCM IP to the CDT1 input levels. Relative CDT1 levels were then quantified in the MCM IP +CST or +CS compared to MCM IP samples without CST expression (Figure 4B and EV4, numbers below blot). Here, we observed a significant decrease in CDT1 binding to MCM in the presence of CST. Interestingly, when only CTC1 and STN1 were expressed, the decrease in CDT1 binding to MCM, while still significant, was not as severe as expression of the entire CST complex. This indicates that binding of TEN1, which likely alters the confirmation of CTC1 and STN1, also contributes to the inhibition of CDT1 binding to MCM. These striking findings indicate that CST disrupts the interaction between MCM and CDT1. We propose that this occurs through CST blocking the CDT1 interaction site on MCM, which prevents/obstructs stable binding of CDT1 (Figure 4C). Disruption of the MCM-CDT1 interaction would directly affect origin licensing (i.e. MCM loading) by preventing MCM recruitment, thus explaining why CST decreases origin licensing.

### STN1 chromatin association increases in S-phase

While examining the levels of chromatin-bound MCM, we also determined whether CST chromatin association changed throughout the cell cycle. HeLa cells stably expressing Flag-STN1 were synchronized by double-thymidine block and collected 0, 1.5, 3, 6, 9, 12 and 24h after release. Flow cytometry was used to verify cell synchronization and EdU incorporation to identify cells in S-phase (Figure 5A). Western blot analysis for STN1 was then performed on chromatin fractions (Figure 5B). At the 1.5-6 h time points, cells were predominately in S-phase and transitioned to G2/M by 9 h. While a small fraction of STN1 remains chromatin-bound throughout the cell cycle, STN1 levels significantly increased on the chromatin at the 1.5-6 hr timepoints (Figure 5B), suggesting that CST is recruited to the chromatin during S-phase. Significantly, the timing of this increase differs from the timing of increased CST telomere association, which occurs in late S/G2 (Chen et al., 2012).

**Figure 5.**
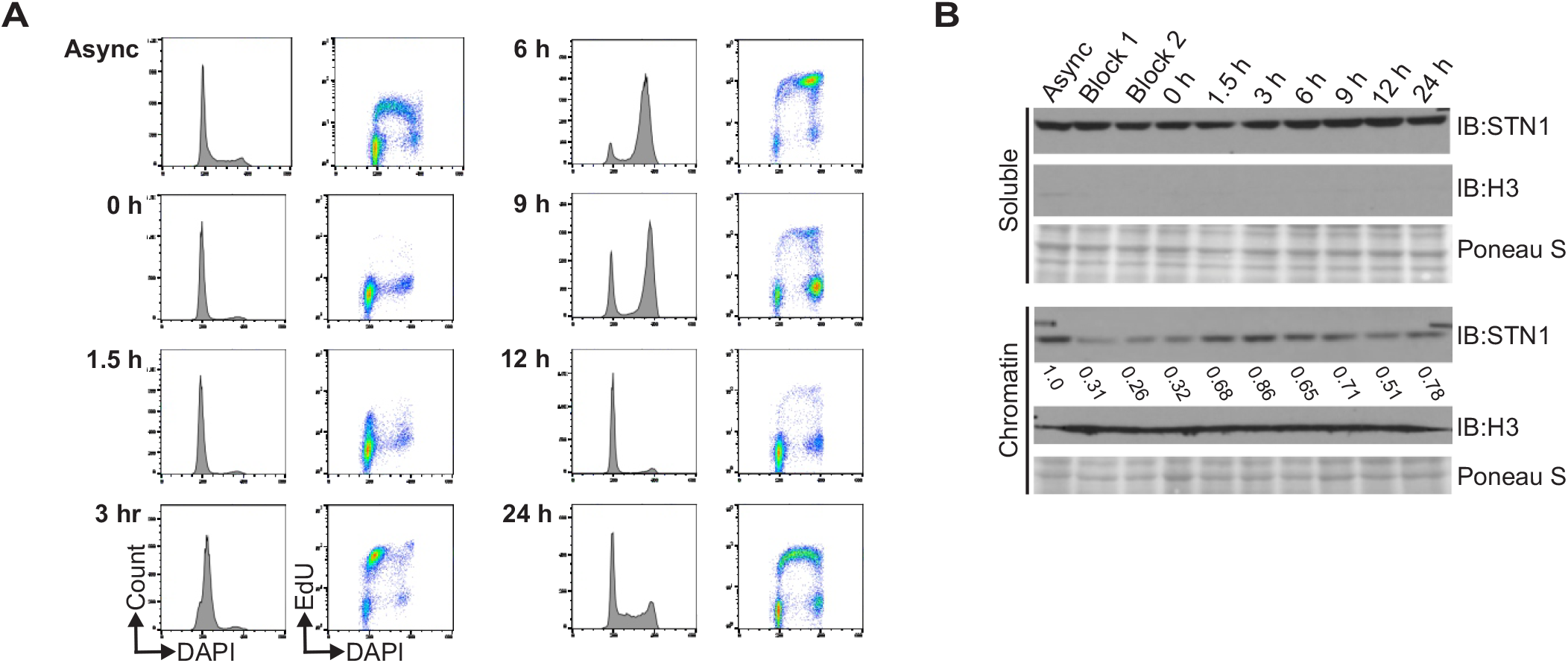
STN1 chromatin association increases in S-phase. (**A**) Cell synchronization by double-thymidine block was confirmed by flow cytometry. Graphs on the left of each timepoint show the DNA content (DAPI) versus number of cells (count) and on the right is the DNA content versus EdU signal intensity. (**B**) Western blot analysis showing soluble and chromatin-associated STN1 throughout the cell cycle. Async: asynchronous cells; Block 1: cells after first thymidine block; Block 2: cells prior to second thymidine block. Ponceau S stains total protein and was used as a loading control. Histone H3 was used as a control for chromatin fractionation. Numbers below the STN1 chromatin blot indicate the relative levels of STN1 throughout the cell cycle compared to the asynchronous control. Data is representative of three independent biological experiments.

### CST interacts with AND-1 and promotes AND-1 and pol α chromatin binding

Previous studies have shown that CST interacts with pol α and depletion of CST subunits leads to anaphase bridges and chromosome fragility in the absence of exogenous replication stress (Chastain et al., 2016, Chen, Majerska et al., 2013, Goulian et al., 1990, Stewart et al., 2012, Wang et al., 2012, Wang & Chai, 2018). These observations suggest CST plays additional roles in DNA replication beyond origin licensing. Given that CST is recruited to the chromatin in S-phase and interacts with pol α, we wondered whether CST promotes replisome assembly and origin firing, particularly whether CST affects the association of AND-1 and pol α with the replisome.

Prior to origin firing, pol α is coupled to the replisome by AND-1 (Simon, Zhou et al., 2014). Since CST also physically and functionally interacts pol α, we hypothesized that CST may assist or replace AND-1 to link pol α to the replisome under certain situations, such as at G4s or other GC-rich DNA, which are enriched at origins. AND-1, also known as Ctf4 and WDHD1, is a homotrimeric complex composed of conserved WD40 repeats and a SepB domain (Guan, Li et al., 2017). Human AND-1 also contains a HMG domain, which facilitates its interaction with pol α, and was recently shown to bind to ssDNA *in vitro*, which may help to it position pol α for primer synthesis (Kilkenny, Simon et al., 2017).

We first tested whether depletion of STN1 altered chromatin-bound AND-1 in HeLa cells. The cells were pre-extracted and chromatin-associated AND1 was measured by IF (Figure 6A-B). STN1 knockdown caused a significant decrease in AND-1 levels, suggesting that CST promotes AND-1 chromatin binding. To ensure that CST had not decreased total cellular AND-1, we checked AND-1 levels in whole cell lysates, where AND-1 was slightly increased in the shSTN1 and shSTN1-Res compared to shNT cells (Figure EV5). While the reason for these changes are not clear, they indicate that decreased AND-1 chromatin association is not due to decreased cellular AND-1 levels. We also examined AND-1 and pol α in HCT116 shSTN1 cells and found a decrease in both AND-1 and pol α chromatin association (Figure 6C). These findings fit with a role for CST in replisome assembly, as AND-1 association is needed for pre-IC formation. However, it is interesting to note that only a fraction of AND-1 is lost from the chromatin, suggesting that CST is only required at subset of replication origins to recruit AND-1.

**Figure 6.**
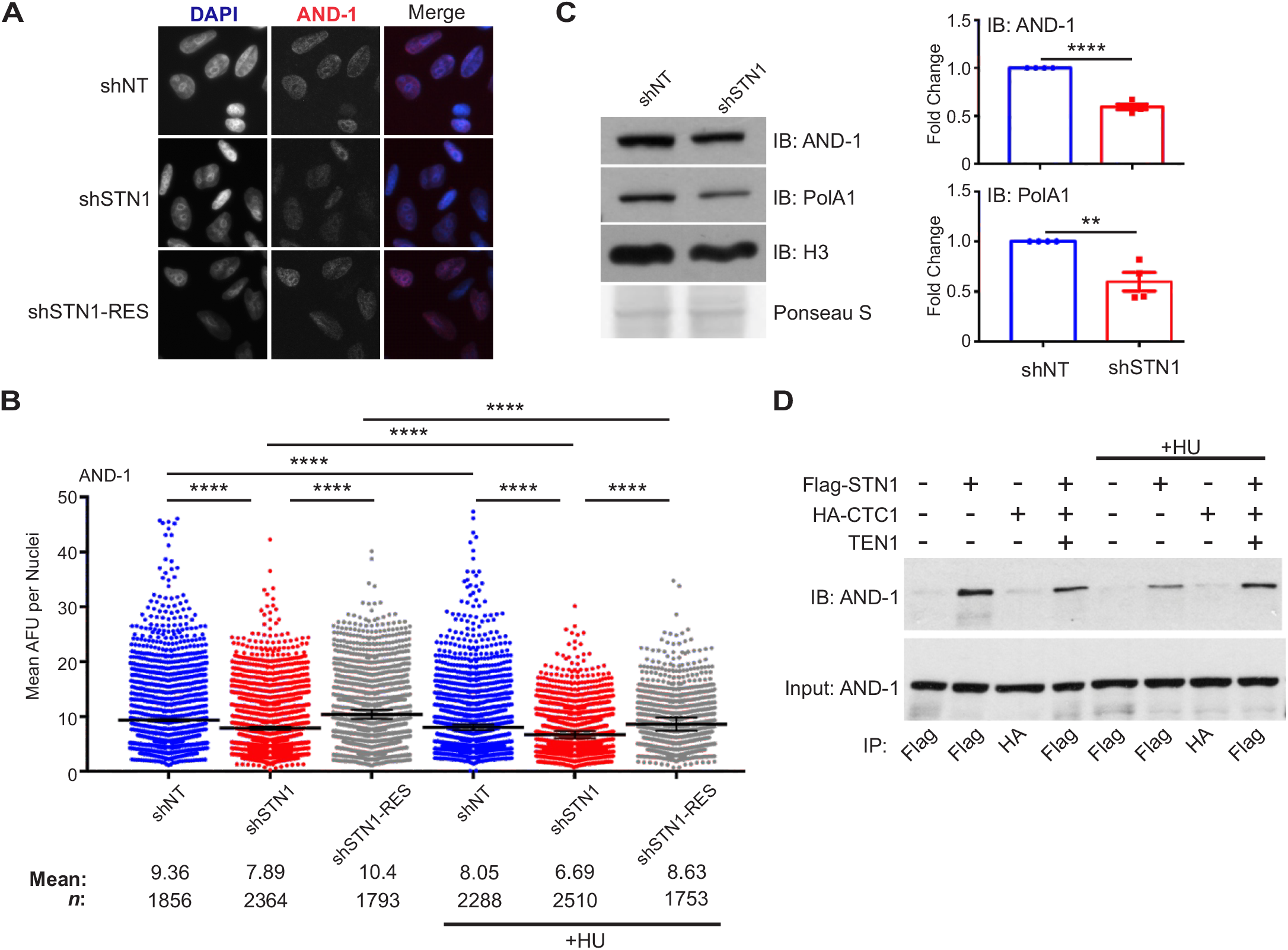
CST is required for AND-1 and pol α chromatin association. (**A**) Representative images of pre-extracted HeLa cells used to measure chromatin-associated AND-1. DAPI: blue, AND-1: red. (**B**) Dot plots of mean AND-1 intensity per nuclei in AFU for each cell line, as indicated. Black line and numbers below the graph indicate the mean AFU. Error bars indicate the +/-SEM of three independent biological experiments. (**C**) Western blot analysis showing chromatin fractions from HCT116 cells. Ponceau S stains total protein and was used a loading control. AND-1 and pol α levels were normalized to shNT in the graph. (**D**) Co-IP was performed with Flag or HA antibody in cell lysates from HEK 293T cells, as indicated. 5% input was loaded as a control. For +HU samples in (B) and (D), HU was added for 2 h prior to collection. Data is representative of three independent biological experiments. *n*=indicates the number of total nuclei scored. *P*-values were calculated by an unpaired, two-tailed Mann-Whitney test in (B) and student’s t-test in (C) (\*\*\*\**p≤*0.0001, ** *p≤*0.01).

To address the possibility that CST indirectly affects AND-1 levels, we next determined whether AND-1 and CST physically associate. Co-IP was performed in cells expressing Flag-STN1, HA-CTC1 or the entire CST complex. We found that STN1 alone or in complex with CST interact with AND-1 (Figure 6D). These results provide evidence that CST directly aids in the loading of AND-1/pol α on the chromatin and hence with the replisome.

Given our previous finding that CST promotes dormant origin firing in response to HU treatment, we next asked whether the dependence of AND-1 on CST for its chromatin association could explain how CST promotes dormant origin firing in response to genome-wide replication fork stalling (Stewart et al., 2012, Wang et al., 2014). Although HU inhibits global origin firing, it promotes the firing of dormant replication origins nearby stalled forks (Ge & Blow, 2010). We hypothesized that CST might enable firing of these dormant origins by recruiting AND-1/pol α. A prediction of this hypothesis is that STN1 knockdown would decrease AND-1 association with dormant origins nearby forks that stalled following HU treatment. If correct, then this would cause an additional decrease in chromatin-bound AND-1 on top of the decrease already observed after STN1 knockdown in the absence of HU treatment. Thus, the effect of STN1 knockdown on AND-1 chromatin association should be much larger in HU-treated cells.

To test whether fork stalling affected AND-1 levels on the chromatin, cells were treated with HU for 2 h and IF for AND-1 performed (Figure 6B). With HU treatment, we observed decreased AND-1 chromatin association across all cell lines, suggesting that HU-induced replication fork stalling causes an overall decrease in AND-1 chromatin association. This overall decrease in AND-1 is likely due to the repression of global origin firing. When STN1 depleted cells were treated with HU, we observed an additional decrease in AND-1 chromatin association relative to control cells. However, the magnitude of the decrease was the same in the HU-treated and untreated cells (∼ 15%). Since the effect of STN1 knockdown was not magnified in HU-treated cells, this finding indicates that CST is unnecessary for AND-1 to associate with the dormant origins that are fired in response to genome-wide replication fork stalling. We therefore infer that CST associates with AND-1 and pol α to stabilize replisome formation at a subset of origins that are active during an unperturbed cell cycle.

## Discussion

In this study, we show that CST regulates two different aspects of DNA replication. First, CST interacts with the MCM complex and disrupts its interaction with CDT1, leading to the suppression of origin licensing. Second, CST associates with AND-1 to promote AND-1 and pol α chromatin association, presumably with the replisome. It is striking that CST functions in two quite separate aspects of DNA replication but this is perhaps not surprising given the RPA-like nature of CST and that RPA also functions in multiple aspects of DNA replication and repair (Fanning et al., 2006). Moreover, CST has already been shown to play distinct roles in several aspects of telomere replication (Stewart et al., 2018). It is interesting that the newly discovered roles of CST in origin licensing and replisome assembly are independent of global replication fork stalling, as this suggests that CST’s roles in dormant origin firing and RAD51 recruitment following genome-wide fork stalling are distinct activities (Chastain et al., 2016, Stewart et al., 2012, Wang & Chai, 2018). At present, the mechanism by which CST facilitates replication restart following genome-wide fork stalling and whether it is directly or indirectly mediated is unclear. However, here we show direct involvement of CST in origin licensing and replisome assembly.

Origin licensing requires CDT1 interaction with MCM to enable MCM binding to ORC/CDC6 and sequential loading of two MCM hexamers on the chromatin (Ticau et al., 2017, Ticau et al., 2015). Our results demonstrate that CST disrupts the interaction between MCM and CDT1 (Figure 4), which explains how CST suppresses origin licensing (Figure 7A). Prior structural analysis of yeast MCM and OCCM complexes, combined with our current data, provide insight into the molecular details whereby CST blocks the MCM-CDT1 interaction and MCM loading (Abid Ali, Douglas et al., 2017, Li, Zhai et al., 2015, Yuan, Bai et al., 2016, Yuan, Riera et al., 2017, Zhai, Cheng et al., 2017a, Zhai et al., 2017b). Our yeast-two-hybrid results indicate that STN1 strongly interacts with the MCM4-MCM7 interface (Figure 3). Within the MCM hexamer, MCM4-MCM7 are located opposite of the MCM2-MCM5 gate used for loading MCM. Recent work suggests that, in yeast, CDT1 interacts with the MCM2-MCM6-MCM4 interface and stabilizes an open ring conformation for MCM loading (Frigola et al., 2017). Additionally, NMR studies showed that the C-terminal domain of MCM6 interacts with CDT1, suggesting a similar binding interface in human cells (Liu et al., 2012, Wei, Liu et al., 2010). Finally, we find that human CDT1 also interacts with MCM4 (Figure 4A).

**Figure 7.**
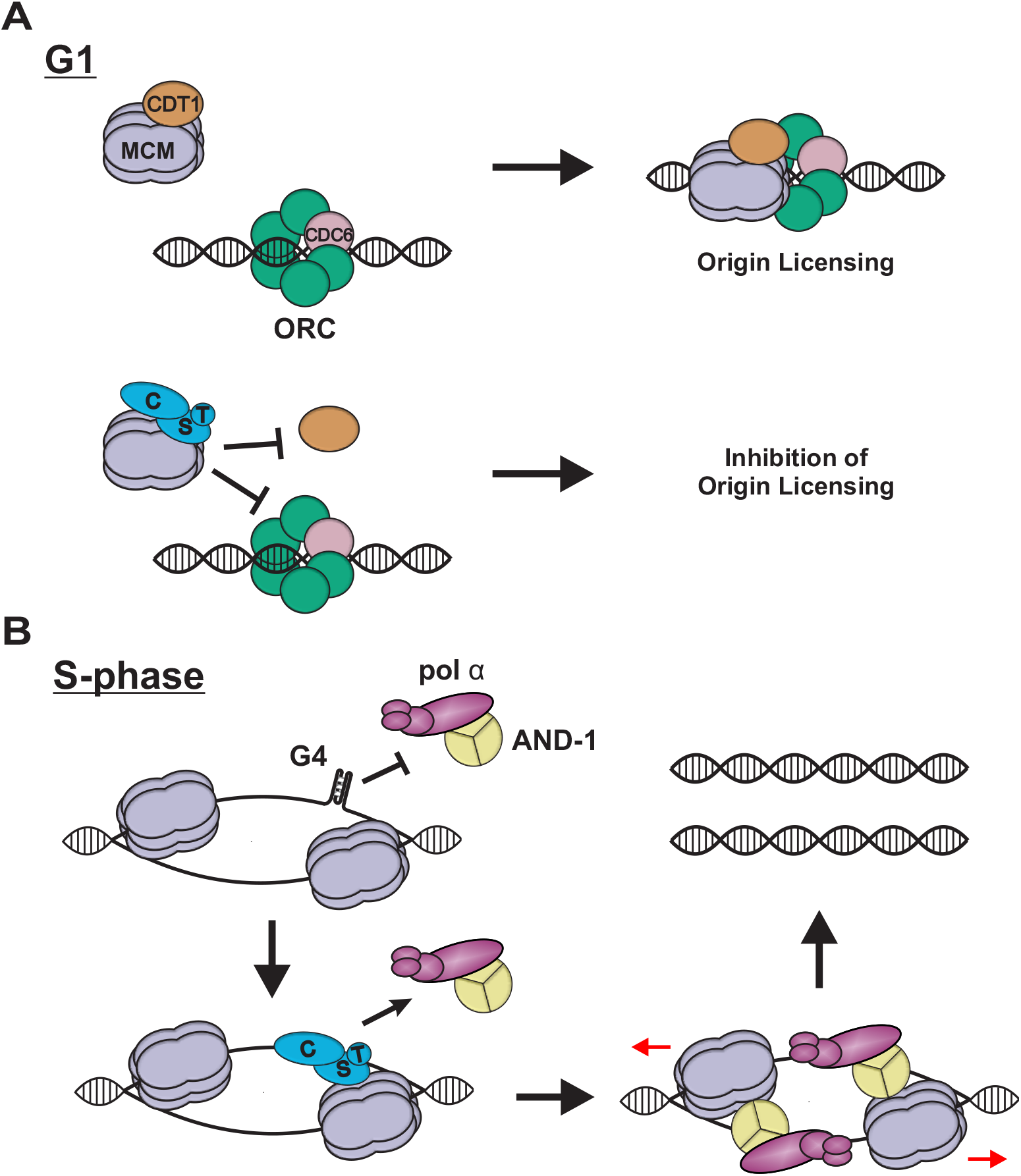
Potential roles of CST during DNA replication. (**A**) In G1, CDT1 interacts with MCM, which facilitates interaction with ORC/CDC6 and origin licensing (top). When CST binds to MCM, this prevent CDT1 from interacting with MCM, blocking origin licensing (bottom). (**B**) During S-phase, CST may prevent or remove DNA secondary structures, such as G4s, that inhibit AND-1 from associating with the replisome, only MCM is shown for simplicity. Following removal of such structures, AND-1 and pol α associate with CST, leading replisome assembly and initiation of DNA synthesis.

Binding of CST to MCM4, which is adjacent to MCM6, could destabilize/block the interaction between CDT1 and MCM (see Figure 3C), leading to a closed ring conformation and the inability for MCM to interact with ORC/CDC6 (Figure 7A). Alternatively, CST binding could cause a conformational shift that prevents CDT1 binding to MCM. Disruption of the MCM-CDT1 interaction may be used to prevent the loading of malformed or incomplete MCM hexamers at origins, or overloading of MCM at specific regions of the genome, which could affect replication timing (Das, Borrman et al., 2015). In budding yeast, CDC6 is known to prevent unproductive MCM loading following OCCM formation (Frigola, Remus et al., 2013). CST could provide an additional quality control mechanism to prevent unproductive MCM loading prior OCCM formation. However, further studies are needed to determine the precise role of CST in preventing origin licensing.

With regard to the timing of CST association with MCM, studies to detect replisome-associated proteins have failed to detect CST at replisomes actively synthesizing DNA (Miyake et al., 2009, Sirbu et al., 2013). However, our data suggest that CST interacts with CMG prior to origin firing. These data suggest that CST interacts with MCM during pre-RC/replisome formation and may not be required during normal DNA synthesis. Instead, we propose that CST is recruited at regions of difficult-to-replicate G-rich DNA to promote replication restart and replication initiation by recruiting AND-1/pol α (Stewart et al., 2012) (Figure 7).

Altered origin licensing can have a profound impact on genome stability, particularly when combined with defects in other pre-RC or pre-IC factors or oncogene-induced replication stress (Kotsantis et al., 2018). For example, mutations in pre-RC factors, which decrease origin licensing, are known to cause Meier-Gorlin syndrome, a primordial dwarfism disorder (Bicknell, Bongers et al., 2011, Bicknell, Walker et al., 2011). Changes in STN1 mRNA expression were also reported in a number of cancers (Chastain et al., 2016). Thus, future research is required to determine how changes in CST expression or mutations affect origin licensing under different conditions. It will also be essential to understand the extent to which defects in origin licensing are counterbalanced with other CST-related functions, such as dormant origin activation following replication stress.

We also find that CST interacts with AND-1 (Figure 5). AND-1 is important for recruitment of pol α, and may also act as a hub for the localization of other proteins to the replisome (Hao, de Renty et al., 2015, Im, Ki et al., 2009, Yoshizawa-Sugata & Masai, 2009, Zhu, Ukomadu et al., 2007). Additionally, AND-1 has ssDNA-binding activity, which may help position pol α for replication (Kilkenny et al., 2017). A recent report demonstrated that auxin-induced degradation of AND-1, in DT40 cells, caused large stretches of ssDNA, DNA damage and G2 arrest but, surprisingly, AND-1 degradation did not prevent the initiation of DNA synthesis (Abe, Kawasumi et al., 2018). The ability of AND-1 deficient cells to still initiate replication may reflect a role for other factors, such as CST and MCM10, in linking pol α to the replisome. However, without AND-1, uncoupling of pol α from the replisome is likely to be common, leading to ssDNA gaps and incomplete replication (Im et al., 2009, Zhu et al., 2007).

Since G4s are enriched at replication origins (Valton & Prioleau, 2016), we propose that the interaction between CST and AND-1 may be necessary at specific regions of the genome, such as G4s or GC-rich DNA, which could block AND-1 binding. Here, we envision that CST interacts with MCM and removes DNA secondary structures that may form upon initial unwinding but prior to DNA synthesis (Figure 7B) (Bhattacharjee et al., 2017). CST could then recruit/stabilize AND-1 and pol α to the replisome followed by CST dissociation. This role for CST would be akin to the RPA “hand-off” mechanism, where RPA guides the recruitment of specific proteins to the DNA, which then trade places with RPA through sequentially dissociation of its OB-folds (Chen & Wold, 2014, Fanning et al., 2006). AND-1 would then position pol α to initiate DNA synthesis. A similar situation could occur at replication forks stalled by G4s or other secondary structures. Here, CST would resolve the block and then recruit AND-1/pol α to reinitiate DNA synthesis. Since CST interacts with MCM and pol α, it is also possible that CST may substitute for AND-1 under certain conditions, as discussed above. It is formally possible that CST could also assist AND-1 with fork restart, fork protection or DNA damage signaling (Abe et al., 2018, Chen, Liu et al., 2017, Hao et al., 2015, Li, Li et al., 2017). However, these scenarios seem unlikely because HU-induced fork stalling did not influence the interaction between CST and AND-1 or further decrease AND-1 levels with STN1 knockdown (Figure 6).

Defects in the recruitment of AND-1/pol α to the replisome could explain the increased anaphase bridges and chromosome fragility seen after depletion or deletion of CST subunits even though bulk, genome-wide DNA replication (as assayed by EdU uptake) remains unaffected (Chastain et al., 2016, Stewart et al., 2012, Wang et al., 2012, Wang & Chai, 2018). The ∼ 15% decrease in chromatin-bound AND-1 in STN1 depleted cells suggests that CST is required for AND-1 association at only a subset of origins. Replication blocks at these origins (e.g. G-rich regions) could prevent origin firing, leading to anaphase bridges or chromosome fragility without affecting bulk DNA replication. Such a requirement for CST would fit with CST being a specialized, as opposed to, a general replication factor. In this case, binding of CST, rather than RPA, would be advantageous because it could resolve the replication block without activating ATR and a subsequent DNA damage response.

Overall, our findings provide evidence that CST functions in two distinct aspects of genome-wide DNA replication, namely origin licensing and replisome assembly. CST was previously shown to interact with pol α and stimulate its activity. Here, we demonstrate that CST interacts with additional replisome components, MCM and AND-1. Future work to untangle how CST expression affects origin licensing, replisome assembly and replication dynamics will help to clarify the non-telomeric roles of CST. Furthermore, determining whether CTC1 or STN1 mutations, arising in Coats plus patients, affect origin licensing and firing through interactions with AND-1 and MCM may help to decipher the molecular pathogenesis of this disease.

## Materials and methods

### Cell Culture

HeLa 1.2.11 cells were maintained in RPMI 1640 media, HCT116 in McCoy’s 5A media and HEK 293T and HeLa TetOn in DMEM at 37°C with 5% CO_2_. All cell lines were supplemented with 10% Fetal Bovine Serum and 1% Penicillin/Streptomycin. Media for HeLa1.2.11 and HCT116 shRNA knockdown cells also contained puromycin (1 μg/mL, Gibco) to maintain selection (Stewart et al., 2012). Doxycycline (1 μg/mL, Sigma) was added to HeLa TetOn wild type and CST overexpressing (CST-OE) cells, as previously described (Wang et al., 2014). Cell lines were regularly checked for mycoplasma contamination.

### Whole Cell Extraction

Cell pellets were suspended in lysis buffer (20 mM Tris pH 8.0, 100 mM NaCl, 1 mM MgCl_2_, 0.1% Igepal, Protease Inhibitors [1 μg/mL Pepstatin A, 5 μg/mL Leupeptin, 1 μg/mL E64, 2 μg/mL Aprotinin, 5 μg/mL Antipain], Phosphatase Inhibitors [4 mM β-glycerophosphate, 4 mM sodium vanadate, 20 mM sodium fluoride]) and incubated on ice for 15 minutes. The samples were then sonicated 3 times (10 sec on/5 sec off) at 40% amplitude, and rested on ice for 5-10 minutes. Extracts were treated with Benzonase (0.0625 U/μL, EMD Millipore) for one hour on ice followed by centrifugation at 19,000 *x g* at 4°C for 5 minutes. The supernatant was collected and analyzed by Western blot. Protein concentrations were determined with the BCA assay (Thermo Scientific).

### Chromatin Fractionation

Cell pellets were suspended in lysis buffer (10 mM Tris pH 7.5, 100 mM NaCl, 1 mM MgCl_2_, 0.34 M sucrose, 0.1% Triton-X, 1mM DTT, Protease Inhibitors, Phosphatase Inhibitors) and then rotated at 4°C for 30 minutes, before centrifugation at 19,000 *x g* for 20 minutes at 4°C. The supernatant was saved as the cytosolic fraction. The pellet was re-suspended in soluble nuclear lysis buffer (10 mM Tris pH 7.5, 100 mM NaCl, 1 mM MgCl_2_, 0.34 M sucrose, 1mM DTT, Protease Inhibitors, Phosphatase Inhibitors) using one half of the volume of the lysis buffer used above. The samples were incubated at 37°C for 10 minutes and centrifuged at 19,000 *x g*. The supernatant was removed and the pellet re-suspended in RIPA buffer (50 mM Tris-HCl pH 8.0, 150 mM NaCl, 1% Triton X-100, 0.1% SDS, 1 mM EDTA, 1 mM DTT, Protease Inhibitors, Phosphatase Inhibitors) followed by sonication twice (10 sec on/5 sec off) at medium intensity. The extracts were rested on ice for ∼ 5 minutes and treated with Benzonase (0.0625 U/μL, EMD Millipore) for 1 h on ice. The samples were then centrifuged at 19,000 *x g* at 4°C for 5 minutes and the supernatant saved as the chromatin fraction. Protein concentrations were determined with the BCA assay, and samples analyzed by Western blot.

### Antibodies

Primary: MCM3 (Santa Cruz, sc-390480), MCM4 (Bethyl, A300-125A), MCM6 (BD Biosciences, 611622), MCM7 (Santa Cruz, sc-22782), CDC45 (Santa Cruz, sc-55569), Actinin (Santa Cruz, sc-17829), OBFC1 (STN1) (Abcam, ab119263), TEN1 (Kasbek et al., 2013), WHDH1 (AND-1) (Novus, NBP1-89091), PolA1 (Bethyl, A302-850A), α-Tubulin (Sigma-Aldrich, T9026), H3 (Cell Signaling, 9715), HA-tag [(anti-mouse: Cell Signaling, 2367)(anti-rabbit: Cell Signaling, 3724)], Flag-tag [(anti-mouse: Sigma, F1804)(anti-rabbit: Thermo, PA1984B)], Myc-tag [(anti-mouse: EMD Millipore, 05-724)(anti-rabbit: Abcam, ab1906)]. Secondary: Thermo: anti-rabbit-HRP (32460); anti-mouse-HRP (32430), anti-goat-HRP (31402); Molecular Probes: goat anti-mouse AlexaFluor 647 (A21235), goat-anti-rabbit AlexaFluor 647 (A21244), goat-anti-rabbit AlexaFluor 594 (A11037), goat-anti-mouse AlexaFluor 594 (A11032), goat-anti-mouse AlexaFluor 488 (A11029), goat-anti-rabbit AlexaFluor 488 (A11034).

### Western blot analysis

15-30 μg of protein, unless otherwise indicated, was run on SDS-PAGE gels and transferred to a nitrocellulose membrane. All membranes were checked with Ponceau S staining for transfer efficiency and then blocked in 5% non-fat milk in PBS plus 0.1% Tween-20 (PBST) for at least 2 h. Primary antibodies were diluted in 5% non-fat milk-PBST or PBS and incubated overnight at 4°C. Primary antibodies were removed and the membranes washed 3x for 10 minutes each in PBST. Secondary antibodies were diluted in 5% non-fat milk-PBST and incubated at least 2 h at RT. After incubation, the membranes were washed 3x for 10 minutes each in PBST. The blots were then developed with Western Lightning Plus ECL (Perkin Elmer) or ECL Prime (GE Healthcare).

### Immunofluorescence

Cells were plated onto coverslips and allowed to grow to 50-70% confluency. They were incubated with 50 μM EdU for 30 minutes, where indicated. For MCM subunits, soluble proteins were pre-extracted with ice-cold 1x CSK buffer (10 mM HEPES pH 7.4, 0.3 M sucrose, 100 mM NaCl, 3 mM MgCl_2_, Protease Inhibitors, Phosphatase Inhibitors) containing 0.1% Triton X-100 for 2-3 min at RT. Cells were then fixed with ice-cold 100% methanol at −20°C for 10 min. Cells were then blocked in a 2% BSA/1% Fish Gelatin-PBS solution for at least 1 h at RT or overnight at 4°C. Primary antibodies were diluted in 2% BSA/1% Fish Gelatin-PBS at 1:500 and incubated with the coverslips for at least 1 h followed by three PBST washes. Secondary antibodies were incubated at 1:1000 with the coverslips for at least 1 h at RT followed by three PBST washes. For AND-1, cells were pre-extracted and fixed as previously described (Chen et al., 2017). Coverslips were then blocked with 3% BSA in 1x PBS followed by incubation with 1:100 α-WDHD1 (AND1) for 1 h at RT. Following three PBST washes, the coverslips were incubated with 1:1000 goat α-rabbit AlexaFluor 594 for 30 min at RT and then washed three times with PBST. Where indicated, EdU was detected per the manufacturer’s instructions (Life Technologies). All coverslips were then dehydrated using an ethanol series and mounted with FluoroGel (Electron Microscopy Sciences) containing 0.2 ug/mL DAPI. Images were taken under 60x oil immersion on an EVOS FL microscope (Thermo Fisher). The nuclear signal intensity was analyzed in ImageJ as previously described (Stewart et al., 2012).

### Cell Synchronization

Cells were plated into 10 cm dishes at ∼ 5 × 105 cells/mL and allowed to grow overnight. 2 mM thymidine was added and plates were incubated for 14 h at 37°C. After 14 h, media was removed and all cells were washed three times with warm 1x PBS. Fresh media was added and cells were released for 9 h. After 9 h, a second thymidine block was initiated by the addition of 2 mM thymidine. Plates were incubated at 37°C for 16 hr. After 16 h, cells were washed three times with warm 1x PBS, and the media replaced. Samples were then collected at specific timepoints (0, 1.5, 3, 6, 9, 12, 24 h release). After collection, cells were divided for flow cytometry (see below) or fractionation and western analysis (see above).

### Flow Cytometry

After cells were collected, the supernatant was removed, the samples were pre-extracted to remove soluble proteins by the addition of 500 μL of fresh 1x CSK buffer plus 0.1% Triton X-100 and incubated at RT for 5 minutes. Leaving the CSK buffer on the cells, 5 mL of ice-cold 100% methanol was then added to the tubes dropwise with gentle vortexing. Tubes were capped, inverted once, and then placed at −20°C for 10 min. Samples were inverted once during incubation to prevent clumping. 5 mL of filter-sterilized 1% BSA-PBS was then directly added to the CSK buffer/100% methanol mixture, tubes inverted several times and samples were centrifuged at 1000 x *g* for 5 minutes. The supernatant was removed and 5 mL of fresh, filter-sterilized 1% BSA-PBS was added to the resuspended cell pellet. Samples were stored at 4°C.

Cells were then spun down at 1000 x *g* for 5 minutes. The supernatant was removed and 200 μL of MCM6 antibody diluted (1:500) in 1% BSA-PBST was added to the resuspended cell pellet for at least 1 h at RT, with mild vortexing halfway through the incubation. 5 mL of 1% BSA-PBST was then added and samples spun down at 1000 x *g* for 5 minutes and the supernatant removed. A second wash of 5 mL 1% BSA-PBST wash was then performed. Cells were then resuspended and incubated in 250 μL of goat α-mouse AlexaFluor 647 (1:500) in 1% BSA-PBST for at least 1 h at RT, protected from light, with mild vortexing halfway through the incubation. Cells were washed twice as described above with 5 mL of 1% BSA-PBST.

EdU was detected by Click-iT chemistry. Reaction cocktail was made according to the manufacturer’s instructions (Thermo Fisher). 250 μL of the complete Click-iT reaction cocktail was added and the samples incubated at RT for 30 minutes. 5 mL of 1% BSA-PBST was then added to each tube and the samples spun down at 1000 x *g* for 5 minutes, the supernatant was removed and the cells resuspended in the residual liquid. 1 mL of fresh DAPI Staining Solution (200 μL 10% Triton X-100, 20 μL 1 mg/mL DAPI, 200 μL 10 mg/mL RNase, into 20 mL 1% BSA-PBS) was added and the cells were incubated for at least 15 minutes at RT. Samples were spun at 50 x *g* for 30 seconds to remove cell clumps and debris through filter-capped tubes (Corning) and run on a BD LSR II Flow Cytometer in the Microscopy and Flow Cytometry Facility at the University of South Carolina, College of Pharmacy. Measurements and analysis were then performed for chromatin-bound MCM6, EdU and DAPI using FlowJo (FlowJo, LLC). Gates for EdU positive cells were created using control samples lacking EdU (Figure 2B and EV2E) and gates for MCM6 positive cells were selected, using control cells without MCM6 antibody (Figure 2D and EV2A). To separate out G1, S and G2/M populations in Figure 2E and EV2G, the following gating was performed (see Figure EV2A-D). G1 cells were EdU-cells at ∼ 200 DAPI peak (2n). S-phase cells were EdU+ cells in the ∼ 200-400 DAPI range (2n-4n). G2/M were EdU-cells in the ∼ 400 DAPI range (4n), which would omit any cells entering G2 during the 30 min EdU labeling period.

### Co-Immunoprecipitation (co-IP) Assay

For co-IP experiments, cells were grown overnight, such that cells were at 50-70% confluency on the day of transfection. 10 μg of total DNA was mixed with 20 μl Polyethylenimine (10 mg/ml) (Polysciences) for each transfection. Plasmids used for transfection include: pcDNA3.1-Flag-STN1 (Stewart et al., 2012), pcDNA3.1-HA-CTC1 (Surovtseva et al., 2009), pTRE2-TEN1 (Wang et al., 2014), pInducer20-Blast-CDT1-HA (a gift from Jean Cook, Addgene plasmid #109335) and pcDNA3.1 Flag-tagged MCM subunits (ORF cDNA clones from Genscript). After 48 h, cells were collected and resuspended in 500 μl lysis buffer (20 mM Tris pH 8.0, 100 mM NaCl, 1 mM MgCl_2_, 0.1% Igepal) with Protease and Phosphatase inhibitors, incubated on ice for 15 min and then treated with Benzonase for 1 h at 4°C with rotation to digest the DNA. After incubation, samples were spun down at 19,000 *x g* for 7 min at 4°C and the supernatant moved to a new tube. Lysates were pre-cleared by adding 50 μL of Protein A/G bead slurry (Santa Cruz) and incubating at 4°C for 30-60 min with rotation. Beads were pelleted at 1000 *x g* for 1 min at 4°C and the supernatant collected. For IP, 30 μL of 50% M2 α-Flag (Sigma) or α-HA (Sigma) agarose bead slurry was added to 500 ug pre-cleared whole cell lysate in a total volume of 300 μL and incubated overnight at 4°C with rotation. Beads were then pelleted by centrifugation at 1000 *x g* for 1 min at 4°C and were washed four times with 1 mL of lysis buffer at 4°C for 5 min with rotation for each wash. Proteins were released from the beads by adding 75 μL 2x sample buffer (100mM Tris-HCl pH 6.8, 4% SDS, 20% glycerol and 0.02% bromophenol blue). Samples were boiled for 5 mins before SDS-PAGE and Western blot analysis. For quantification of the co-IP results in Figure 4B and EV4, the relative level of MCM pulled down with CST expression was normalized to the samples without CST expression [MCM(+CST)/MCM(-CST)=Relative IP levels]. CDT1 in the input were then normalized to CDT1 input levels [CDT1(IP)/CDT1(input)=CDT1 level]. CDT1 levels were then divided by the relative MCM IP. For example, [(CDT1(IP)/CDT1(input))/(MCM2(+CST) IP/MCM2(-CST) IP)=Relative CDT1 in MCM2 IP]. We also quantified CDT1 association to MCM without normalizing for CDT1 input levels and also observed a significant decrease (>50%) in CDT1 association with MCM subunits with CST expression.

### Yeast Two-Hybrid Screens

The human CTC1, STN1, TEN1, MCM2, MCM3, MCM4, MCM5, MCM6, MCM7 or CDC45 were amplified by PCR using Phusion polymerase (Thermo) from pcDNA3.1 plasmids encoding the cDNA of each protein (MCM plasmids from Genscript). The PCR products were then cloned into the pGBKT7-BD or pGADT7-AD plasmids. NdeI/SalI sites were used to link CTC1, STN1 and TEN1 to pGBKT7-BD while NdeI/XhoI sites were used for pGADT7-AD. For MCM2-7 and CDC45, EcoRI/NdeI sites were used for both plasmids. Mating was performed using *Saccharomyces cerevisiae* strains Y187 and AH109 transformed with pGBKT7 plasmids or pGADT7 plasmids, respectively. The pGADT7-blank or pGBKT7-blank plasmids were used as negative controls. Healthy diploids (only large [2∼ 3 mm], fresh [<2 months old] colonies) on double dropout (DDO) plates, lacking leucine and tryptophan, were selected and cultured in 5 mL YPDA medium overnight at 30°C with shaking at 200 rpm, according to the high-stringency selection protocol. Overnight yeast cultures were diluted to OD600 = 1.0, 0.1, 0.01, from left to right and spotted on DDO, triple dropout (TDO), lacking tryptophan, histidine and leucine, or quadruple dropout (QDO) plates, lacking adenine, histidine, leucine and tryptophan. Incubation was performed at 30°C for up to 10 days. Yeast transformation, mating, interaction test and plasmid isolation were performed using the Yeast Protocols Handbook and Matchmaker GAL4TM Two-hybrid System 3 & Libraries User Manual (Clontech).

### Yeast Protein Extraction

Yeast cells from the selective media plates were cultured in liquid DDO medium overnight at 30°C. 3 mL yeast culture was spun down and the supernatant discarded. The pellet was resuspended with 150 μL of 2 M LiAc, mixed thoroughly and incubated on ice for 5 min. The samples were then spun down at 850 *x g* for 5 min at 4°C and the supernatant discarded. The pellet was then resuspended in 100 μL 2x sample buffer and boiled for 5 min. The samples were spun again and the supernatant transferred to a new tube. The supernatant, containing yeast whole protein, was loaded onto an SDS-PAGE gel and analyzed by Western blot.

## Acknowledgements

We would like to thank Carolyn Price, Michael Wyatt, Alan Waldman, Feng Wang, Anukana Bhattachajree and members of the Stewart lab for critical reading of the manuscript and helpful discussions. We would also like to thank Ji’Vone Freeman and Jazmine Benjamin for assistance performing experiments. This study utilized the services of the Flow Cytometry Core Facility of the COBRE Center for Targeted Therapeutics, supported by NIH grant 5P20GM109091, at the University of South Carolina with assistance from Chang-uk Lim. This work was supported by the National Institutes of Health [R00GM104409] and startup funds from the University of South Carolina to JAS. BC is supported in part by the Magellan Scholar Program through the University of South Carolina.

## Author Contributions

YW, KSB, BC and SMA designed and performed experiments and provided intellectual input. JAS conceived and supervised the study and wrote the manuscript with input from all co-authors.

## References

Abe T, Kawasumi R, Giannattasio M, Dusi S, Yoshimoto Y, Miyata K, Umemura K, Hirota K, Branzei D (2018) AND-1 fork protection function prevents fork resection and is essential for proliferation. Nat Commun 9: 3091

Abid Ali F, Douglas ME, Locke J, Pye VE, Nans A, Diffley JFX, Costa A (2017) Cryo-EM structure of a licensed DNA replication origin. Nat Commun 8: 2241

Anderson BH, Kasher PR, Mayer J, Szynkiewicz M, Jenkinson EM, Bhaskar SS, Urquhart JE, Daly SB, Dickerson JE, O’Sullivan J, Leibundgut EO, Muter J, Abdel-Salem GM, Babul-Hirji R, Baxter P, Berger A, Bonafe L, Brunstom-Hernandez JE, Buckard JA, Chitayat D et al. (2012) Mutations in CTC1, encoding conserved telomere maintenance component 1, cause Coats plus. Nat Genet 44: 338–42

Armanios M, Blackburn EH (2012) The telomere syndromes. Nat Rev Genet 13: 693–704

Barazas M, Annunziato S, Pettitt SJ, de Krijger I, Ghezraoui H, Roobol SJ, Lutz C, Frankum J, Song FF, Brough R, Evers B, Gogola E, Bhin J, van de Ven M, van Gent DC, Jacobs JJL, Chapman R, Lord CJ, Jonkers J, Rottenberg S (2018) The CST Complex Mediates End Protection at Double-Strand Breaks and Promotes PARP Inhibitor Sensitivity in BRCA1-Deficient Cells. Cell Rep 23: 2107–2118

Bhattacharjee A, Wang Y, Diao J, Price CM (2017) Dynamic DNA binding, junction recognition and G4 melting activity underlie the telomeric and genome-wide roles of human CST. Nucleic Acids Res 45: 12311–12324

Bicknell LS, Bongers EM, Leitch A, Brown S, Schoots J, Harley ME, Aftimos S, Al-Aama JY, Bober M, Brown PA, van Bokhoven H, Dean J, Edrees AY, Feingold M, Fryer A, Hoefsloot LH, Kau N, Knoers NV, Mackenzie J, Opitz JM et al. (2011) Mutations in the pre-replication complex cause Meier-Gorlin syndrome. Nat Genet 43: 356–9

Bicknell LS, Walker S, Klingseisen A, Stiff T, Leitch A, Kerzendorfer C, Martin CA, Yeyati P, Al Sanna N, Bober M, Johnson D, Wise C, Jackson AP, O’Driscoll M, Jeggo PA (2011) Mutations in ORC1, encoding the largest subunit of the origin recognition complex, cause microcephalic primordial dwarfism resembling Meier-Gorlin syndrome. Nat Genet 43: 350–5

Briggs TA, Abdel-Salam GM, Balicki M, Baxter P, Bertini E, Bishop N, Browne BH, Chitayat D, Chong WK, Eid MM, Halliday W, Hughes I, Klusmann-Koy A, Kurian M, Nischal KK, Rice GI, Stephenson JB, Surtees R, Talbot JF, Tehrani NN et al. (2008) Cerebroretinal microangiopathy with calcifications and cysts (CRMCC). Am J Med Genet A 146A: 182–90

Casteel DE, Zhuang S, Zeng Y, Perrino FW, Boss GR, Goulian M, Pilz RB (2009) A DNA polymerase-{alpha}{middle dot}primase cofactor with homology to replication protein A-32 regulates DNA replication in mammalian cells. J Biol Chem 284: 5807–18

Cayrou C, Ballester B, Peiffer I, Fenouil R, Coulombe P, Andrau JC, van Helden J, Mechali M (2015) The chromatin environment shapes DNA replication origin organization and defines origin classes. Genome Res 25: 1873–85

Chastain M, Zhou Q, Shiva O, Fadri-Moskwik M, Whitmore L, Jia P, Dai X, Huang C, Ye P, Chai W (2016) Human CST Facilitates Genome-wide RAD51 Recruitment to GC-Rich Repetitive Sequences in Response to Replication Stress. Cell Rep 16: 2048

Chen LY, Majerska J, Lingner J (2013) Molecular basis of telomere syndrome caused by CTC1 mutations. Genes Dev 27: 2099–108

Chen LY, Redon S, Lingner J (2012) The human CST complex is a terminator of telomerase activity. Nature 488: 540–4

Chen R, Wold MS (2014) Replication protein A: single-stranded DNA’s first responder: dynamic DNA-interactions allow replication protein A to direct single-strand DNA intermediates into different pathways for synthesis or repair. Bioessays 36: 1156–61

Chen Y, Liu H, Zhang H, Sun C, Hu Z, Tian Q, Peng C, Jiang P, Hua H, Li X, Pei H (2017) And-1 coordinates with CtIP for efficient homologous recombination and DNA damage checkpoint maintenance. Nucleic Acids Res 45: 2516–2530

Das SP, Borrman T, Liu VW, Yang SC, Bechhoefer J, Rhind N (2015) Replication timing is regulated by the number of MCMs loaded at origins. Genome Res 25: 1886–92

Deegan TD, Diffley JF (2016) MCM: one ring to rule them all. Curr Opin Struct Biol 37: 145–51

Dewar JM, Budzowska M, Walter JC (2015) The mechanism of DNA replication termination in vertebrates. Nature 525: 345–50

Dewar JM, Low E, Mann M, Raschle M, Walter JC (2017) CRL2(Lrr1) promotes unloading of the vertebrate replisome from chromatin during replication termination. Genes Dev 31: 275–290

Fanning E, Klimovich V, Nager AR (2006) A dynamic model for replication protein A (RPA) function in DNA processing pathways. Nucleic Acids Res 34: 4126–37

Feng X, Hsu SJ, Bhattacharjee A, Wang Y, Diao J, Price CM (2018) CTC1-STN1 terminates telomerase while STN1-TEN1 enables C-strand synthesis during telomere replication in colon cancer cells. Nat Commun 9: 2827

Feng X, Hsu SJ, Kasbek C, Chaiken M, Price CM (2017) CTC1-mediated C-strand fill-in is an essential step in telomere length maintenance. Nucleic Acids Res 45: 4281–4293

Frigola J, He J, Kinkelin K, Pye VE, Renault L, Douglas ME, Remus D, Cherepanov P, Costa A, Diffley JFX (2017) Cdt1 stabilizes an open MCM ring for helicase loading. Nat Commun 8: 15720

Frigola J, Remus D, Mehanna A, Diffley JF (2013) ATPase-dependent quality control of DNA replication origin licensing. Nature 495: 339–43

Ganduri S, Lue NF (2017) STN1-POLA2 interaction provides a basis for primase-pol alpha stimulation by human STN1. Nucleic Acids Res 45: 9455–9466

Ge XQ, Blow JJ (2010) Chk1 inhibits replication factory activation but allows dormant origin firing in existing factories. J Cell Biol 191: 1285–97

Goulian M, Heard CJ, Grimm SL (1990) Purification and properties of an accessory protein for DNA polymerase alpha/primase. J Biol Chem 265: 13221–30

Gu P, Chang S (2013) Functional characterization of human CTC1 mutations reveals novel mechanisms responsible for the pathogenesis of the telomere disease Coats plus. Aging Cell 12: 1100–9

Gu P, Min JN, Wang Y, Huang C, Peng T, Chai W, Chang S (2012) CTC1 deletion results in defective telomere replication, leading to catastrophic telomere loss and stem cell exhaustion. EMBO J 31: 2309–21

Guan C, Li J, Sun D, Liu Y, Liang H (2017) The structure and polymerase-recognition mechanism of the crucial adaptor protein AND-1 in the human replisome. J Biol Chem 292: 9627–9636

Hao J, de Renty C, Li Y, Xiao H, Kemp MG, Han Z, DePamphilis ML, Zhu W (2015) And-1 coordinates with Claspin for efficient Chk1 activation in response to replication stress. EMBO J 34: 2096–110

Higa M, Fujita M, Yoshida K (2017) DNA Replication Origins and Fork Progression at Mammalian Telomeres. Genes (Basel) 8

Im JS, Ki SH, Farina A, Jung DS, Hurwitz J, Lee JK (2009) Assembly of the Cdc45-Mcm2-7-GINS complex in human cells requires the Ctf4/And-1, RecQL4, and Mcm10 proteins. Proc Natl Acad Sci U S A 106: 15628–32

Kasbek C, Wang F, Price CM (2013) Human TEN1 maintains telomere integrity and functions in genome-wide replication restart. J Biol Chem 288: 30139–50

Keller RB, Gagne KE, Usmani GN, Asdourian GK, Williams DA, Hofmann I, Agarwal S (2012) CTC1 Mutations in a patient with dyskeratosis congenita. Pediatr Blood Cancer 59: 311–4

Kilkenny ML, Simon AC, Mainwaring J, Wirthensohn D, Holzer S, Pellegrini L (2017) The human CTF4-orthologue AND-1 interacts with DNA polymerase alpha/primase via its unique C-terminal HMG box. Open Biol 7

Kotsantis P, Petermann E, Boulton SJ (2018) Mechanisms of Oncogene-Induced Replication Stress: Jigsaw Falling into Place. Cancer Discov 8: 537–555

Li N, Zhai Y, Zhang Y, Li W, Yang M, Lei J, Tye BK, Gao N (2015) Structure of the eukaryotic MCM complex at 3.8 A. Nature 524: 186–91

Li Y, Li Z, Wu R, Han Z, Zhu W (2017) And-1 is required for homologous recombination repair by regulating DNA end resection. Nucleic Acids Res 45: 2531–2545

Liu C, Wu R, Zhou B, Wang J, Wei Z, Tye BK, Liang C, Zhu G (2012) Structural insights into the Cdt1-mediated MCM2-7 chromatin loading. Nucleic Acids Res 40: 3208–17

Maizels N (2015) G4-associated human diseases. EMBO Rep 16: 910–22

Masai H, Matsumoto S, You Z, Yoshizawa-Sugata N, Oda M (2010) Eukaryotic chromosome DNA replication: where, when, and how? Annu Rev Biochem 79: 89–130

Matson JP, Dumitru R, Coryell P, Baxley RM, Chen W, Twaroski K, Webber BR, Tolar J, Bielinsky AK, Purvis JE, Cook JG (2017) Rapid DNA replication origin licensing protects stem cell pluripotency. Elife 6

Mirman Z, Lottersberger F, Takai H, Kibe T, Gong Y, Takai K, Bianchi A, Zimmermann M, Durocher D, de Lange T (2018) 53BP1-RIF1-shieldin counteracts DSB resection through CST-and Polalpha-dependent fill-in. Nature 560: 112–116

Miyake Y, Nakamura M, Nabetani A, Shimamura S, Tamura M, Yonehara S, Saito M, Ishikawa F (2009) RPA-like mammalian Ctc1-Stn1-Ten1 complex binds to single-stranded DNA and protects telomeres independently of the Pot1 pathway. Mol Cell 36: 193–206

Moreno SP, Bailey R, Campion N, Herron S, Gambus A (2014) Polyubiquitylation drives replisome disassembly at the termination of DNA replication. Science 346: 477–81

Nakaoka H, Nishiyama A, Saito M, Ishikawa F (2012) Xenopus laevis Ctc1-Stn1-Ten1 (xCST) protein complex is involved in priming DNA synthesis on single-stranded DNA template in Xenopus egg extract. J Biol Chem 287: 619–27

Polvi A, Linnankivi T, Kivela T, Herva R, Keating JP, Makitie O, Pareyson D, Vainionpaa L, Lahtinen J, Hovatta I, Pihko H, Lehesjoki AE (2012) Mutations in CTC1, encoding the CTS telomere maintenance complex component 1, cause cerebroretinal microangiopathy with calcifications and cysts. Am J Hum Genet 90: 540–9

Pozo PN, Cook JG (2016) Regulation and Function of Cdt1; A Key Factor in Cell Proliferation and Genome Stability. Genes (Basel) 8

Rhodes D, Lipps HJ (2015) G-quadruplexes and their regulatory roles in biology. Nucleic Acids Res 43: 8627–37

Riera A, Barbon M, Noguchi Y, Reuter LM, Schneider S, Speck C (2017) From structure to mechanism-understanding initiation of DNA replication. Genes Dev 31: 1073–1088

Romaniello R, Arrigoni F, Citterio A, Tonelli A, Sforzini C, Rizzari C, Pessina M, Triulzi F, Bassi MT, Borgatti R (2013) Cerebroretinal microangiopathy with calcifications and cysts associated with CTC1 and NDP mutations. J Child Neurol 28: 1702–8

Simon AC, Zhou JC, Perera RL, van Deursen F, Evrin C, Ivanova ME, Kilkenny ML, Renault L, Kjaer S, Matak-Vinkovic D, Labib K, Costa A, Pellegrini L (2014) A Ctf4 trimer couples the CMG helicase to DNA polymerase alpha in the eukaryotic replisome. Nature 510: 293–297

Simon AJ, Lev A, Zhang Y, Weiss B, Rylova A, Eyal E, Kol N, Barel O, Cesarkas K, Soudack M, Greenberg-Kushnir N, Rhodes M, Wiest DL, Schiby G, Barshack I, Katz S, Pras E, Poran H, Reznik-Wolf H, Ribakovsky E et al. (2016) Mutations in STN1 cause Coats plus syndrome and are associated with genomic and telomere defects. J Exp Med 213: 1429–40

Sirbu BM, McDonald WH, Dungrawala H, Badu-Nkansah A, Kavanaugh GM, Chen Y, Tabb DL, Cortez D (2013) Identification of proteins at active, stalled, and collapsed replication forks using isolation of proteins on nascent DNA (iPOND) coupled with mass spectrometry. J Biol Chem 288: 31458–67

Sonneville R, Moreno SP, Knebel A, Johnson C, Hastie CJ, Gartner A, Gambus A, Labib K (2017) CUL-2(LRR-1) and UBXN-3 drive replisome disassembly during DNA replication termination and mitosis. Nat Cell Biol 19: 468–479

Stanley SE, Armanios M (2015) The short and long telomere syndromes: paired paradigms for molecular medicine. Curr Opin Genet Dev 33: 1–9

Stewart JA, Wang F, Chaiken MF, Kasbek C, Chastain PD, 2nd, Wright WE, Price CM (2012) Human CST promotes telomere duplex replication and general replication restart after fork stalling. EMBO J 31: 3537–49

Stewart JA, Wang Y, Ackerson SM, Schuck PL (2018) Emerging roles of CST in maintaining genome stability and human disease. Front Biosci (Landmark Ed) 23: 1564–1586

Surovtseva YV, Churikov D, Boltz KA, Song X, Lamb JC, Warrington R, Leehy K, Heacock M, Price CM, Shippen DE (2009) Conserved telomere maintenance component 1 interacts with STN1 and maintains chromosome ends in higher eukaryotes. Mol Cell 36: 207–18

Ticau S, Friedman LJ, Champasa K, Correa IR, Jr., Gelles J, Bell SP (2017) Mechanism and timing of Mcm2-7 ring closure during DNA replication origin licensing. Nat Struct Mol Biol 24: 309–315

Ticau S, Friedman LJ, Ivica NA, Gelles J, Bell SP (2015) Single-molecule studies of origin licensing reveal mechanisms ensuring bidirectional helicase loading. Cell 161: 513–525

Valton AL, Prioleau MN (2016) G-Quadruplexes in DNA Replication: A Problem or a Necessity? Trends Genet 32: 697–706

Wang F, Stewart J, Price CM (2014) Human CST abundance determines recovery from diverse forms of DNA damage and replication stress. Cell Cycle 13: 3488–98

Wang F, Stewart JA, Kasbek C, Zhao Y, Wright WE, Price CM (2012) Human CST has independent functions during telomere duplex replication and C-strand fill-in. Cell Rep 2: 1096– 103

Wang Y, Chai W (2018) Pathogenic CTC1 mutations cause global genome instabilities under replication stress. Nucleic Acids Res 46: 3981–3992

Wei Z, Liu C, Wu X, Xu N, Zhou B, Liang C, Zhu G (2010) Characterization and structure determination of the Cdt1 binding domain of human minichromosome maintenance (Mcm) 6. J Biol Chem 285: 12469–73

Yeeles JT, Deegan TD, Janska A, Early A, Diffley JF (2015) Regulated eukaryotic DNA replication origin firing with purified proteins. Nature 519: 431–5

Yoshizawa-Sugata N, Masai H (2009) Roles of human AND-1 in chromosome transactions in S phase. J Biol Chem 284: 20718–28

Yuan Z, Bai L, Sun J, Georgescu R, Liu J, O’Donnell ME, Li H (2016) Structure of the eukaryotic replicative CMG helicase suggests a pumpjack motion for translocation. Nat Struct Mol Biol 23: 217–24

Yuan Z, Riera A, Bai L, Sun J, Nandi S, Spanos C, Chen ZA, Barbon M, Rappsilber J, Stillman B, Speck C, Li H (2017) Structural basis of Mcm2-7 replicative helicase loading by ORC-Cdc6 and Cdt1. Nat Struct Mol Biol 24: 316–324

Zhai Y, Cheng E, Wu H, Li N, Yung PY, Gao N, Tye BK (2017a) Open-ringed structure of the Cdt1-Mcm2-7 complex as a precursor of the MCM double hexamer. Nat Struct Mol Biol 24: 300– 308

Zhai Y, Li N, Jiang H, Huang X, Gao N, Tye BK (2017b) Unique Roles of the Non-identical MCM Subunits in DNA Replication Licensing. Mol Cell 67: 168–179

Zhu W, Ukomadu C, Jha S, Senga T, Dhar SK, Wohlschlegel JA, Nutt LK, Kornbluth S, Dutta A (2007) Mcm10 and And-1/CTF4 recruit DNA polymerase alpha to chromatin for initiation of DNA replication. Genes Dev 21: 2288–99

