## Supplemental Files for "Human CST suppresses origin licensing and promotes AND-1/Ctf4 chromatin association"

**EXPANDED VIEW FIGURES**

Affiliations:

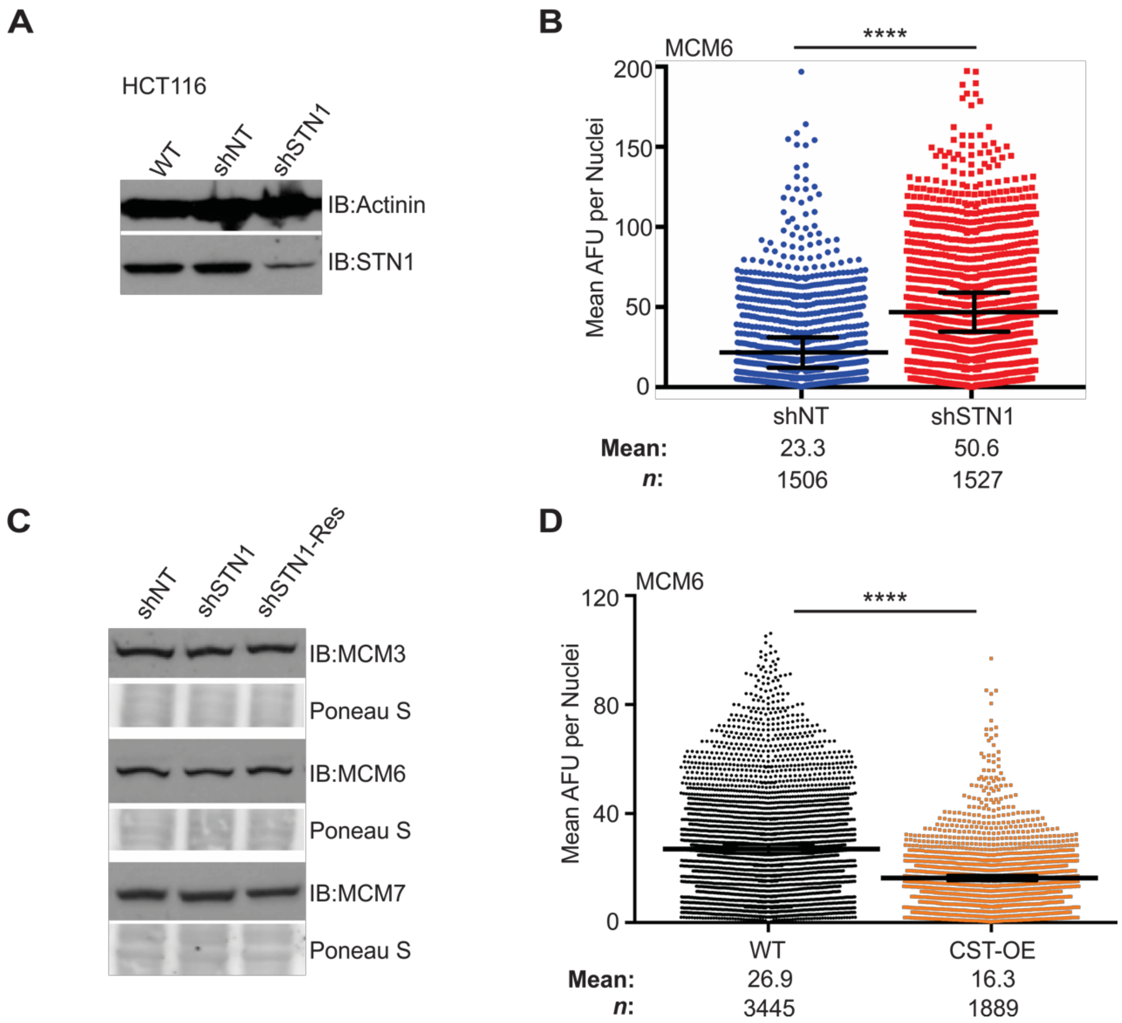


**Figure EV1.** (**A**) Western blot of STN1 levels in HCT116 cell lines. Actinin was used a loading control. (**B**) IF was performed for MCM6 in pre-extracted HCT116 cells. Dot plots of mean MCM6 intensity per nuclei. Black line and numbers below the graph indicate the mean AFU. Error bars represent the +/-SEM of three independent experiments. *n*=indicates the number of total nuclei scored. (**C**) Western blots of MCM levels in HeLa whole cell lysates, as indicated. Ponceau S stain was used as a loading control. (**D**) IF was performed for MCM6 in pre-extracted HeLa cells. Dot plots of mean MCM6 intensity per nuclei in indicated cell lines. Black line and numbers below the graph indicate the mean AFU. Error bars represent the +/-SEM of three independent biological experiments. *n*=indicates the number of total nuclei scored. *P*-values were calculated by an unpaired, two-tailed Mann-Whitney test (****$p \leq 0.0001$).

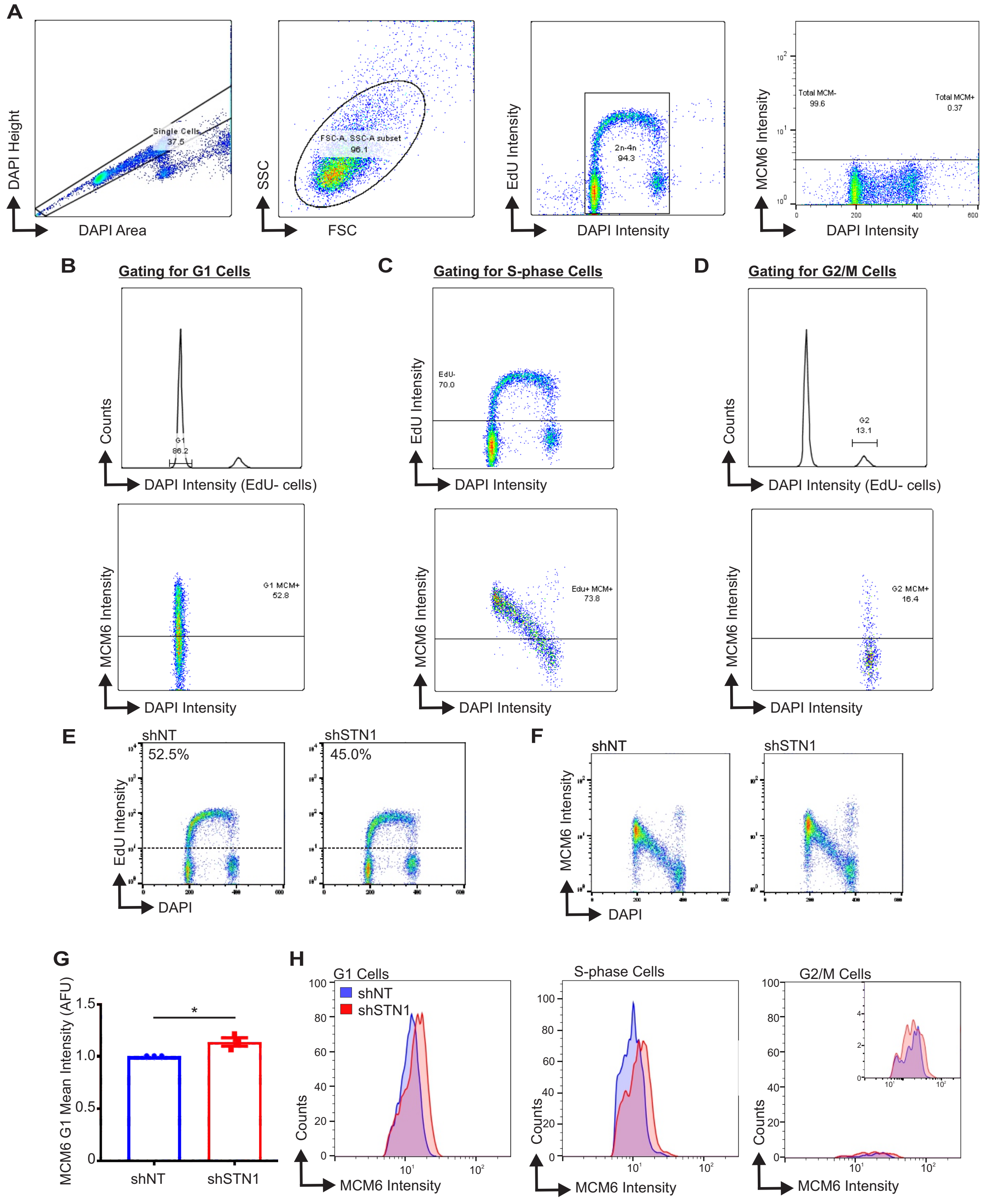


**Figure EV2.** (**A-D**) Gates used for flow cytometry analysis (see Materials and Methods for additional details). (**A**) Gating used for selection of cells for analysis. (**B**) Gating of the G1 MCM+ cells. EdU+ cells were removed from the histogram showing DNA content (top) and the G1 population was then gated. MCM intensity was measured in these G1 cells (bottom). (**C**) Gating of S-phase cells. Top histogram shows the DNA content versus EdU signal intensity and the line denotes the cutoff for EdU+ cells. MCM signal intensity was measured in these EdU+ cells (bottom). (**D**) Gating of the G2/M population. EdU+ cells were removed from the histogram showing DNA content (top). The G2/M population was then gated. MCM signal intensity was measured in these G2/M cells (bottom). Line in the bottom histogram in (B-D) indicates MCM6+ cells, which is based on controls without MCM antibody. (**E-H**) HCT116 cells were treated with EdU, pre-extracted and fixed and flow cytometry performed to detect MCM6, EdU and DAPI. Data is representative of three independent experiments. (**E**) DNA content (DAPI) versus EdU signal intensity. (**F**) DNA content versus MCM6 intensity. (**G**) Graph of the intensity of G1 MCM6 positive cells. Data was normalized to the shNT control. (**G**) Histogram of MCM6 cells (counts) versus their signal intensity in different cell cycle phases, as indicated. *P*-values were calculated by an unpaired, two-tailed student's t-test in (D) (* $p \leq 0.05$).

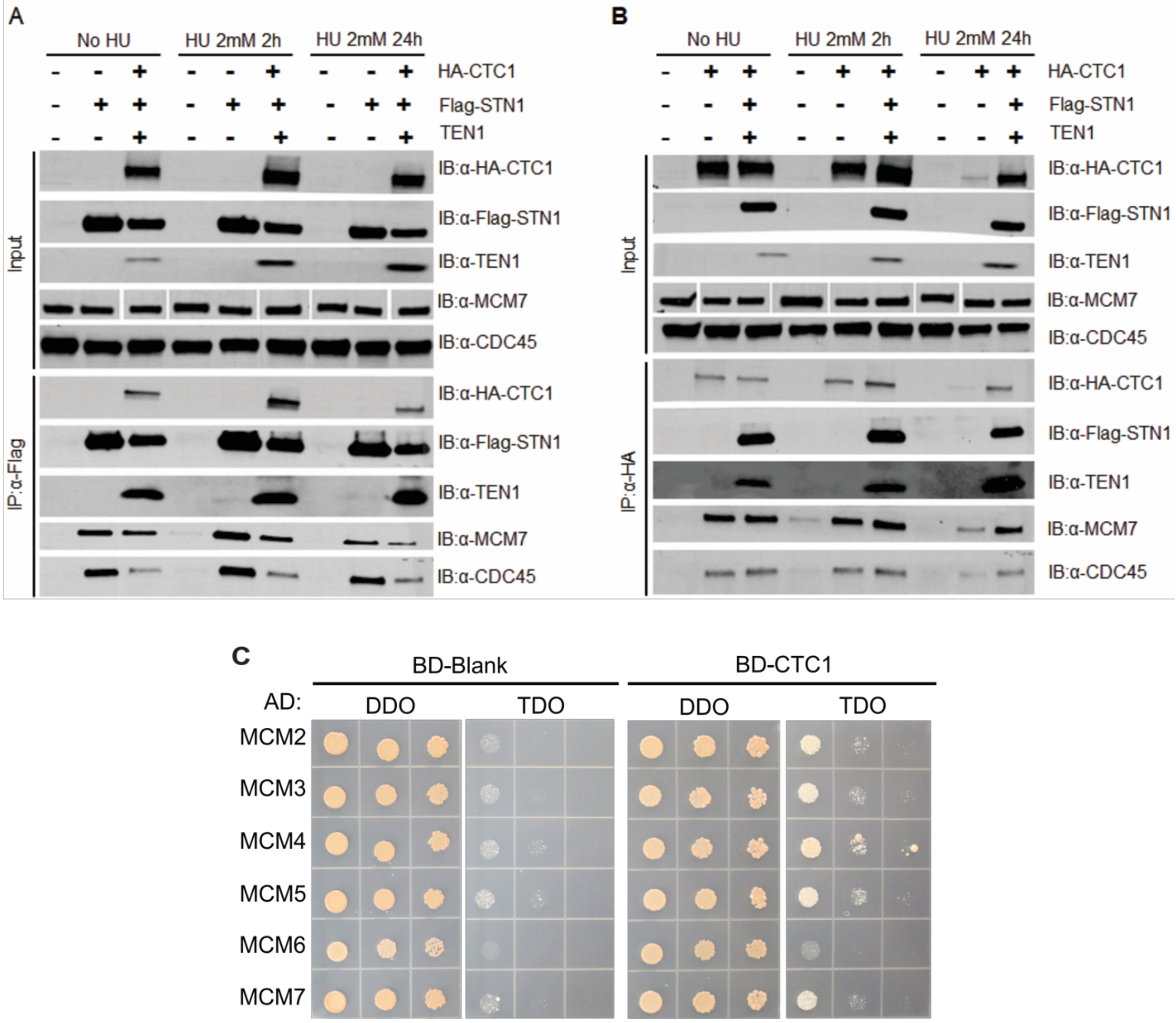


**Figure EV3.** (**A-B**) Co-IP was performed with nuclease-treated lysates from HEK 293T cells expressing Flag-tagged STN1, HA-tagged CTC1 or the CST complex. Indicated samples were treated with HU, as indicated, prior to collection. (**C**) Yeast diploids growing on double synthetic dropout (DDO) medium were selected on triple dropout (TDO) medium. Colony growth was monitored after 4 days of incubation.

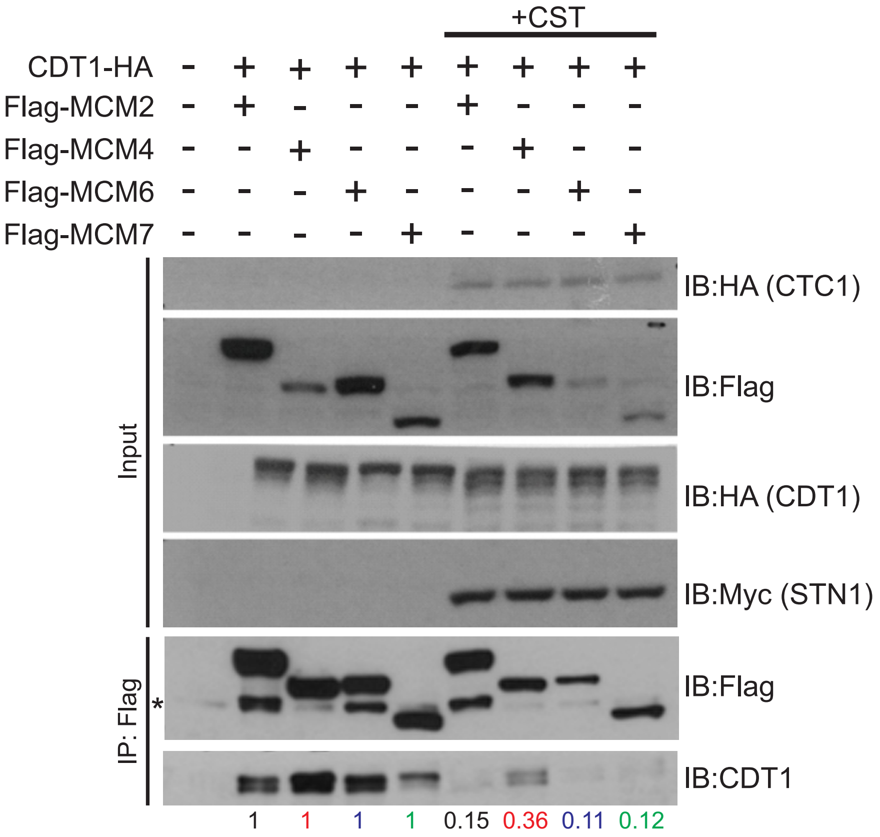


**Figure EV4.** Replicate experiment of Figure 4B. Co-IP assay of lysates from HEK 293T cells expressing Flag-tagged MCM subunits, CDT1-HA together with the CST complex (+CST) or mock plasmids. Lysate from transfected HEK 293T cells were subjected to IP with Flag antibody. Relative CDT1-HA band intensity was then compared +/-CST, as indicated below the gels (see Material and Methods).

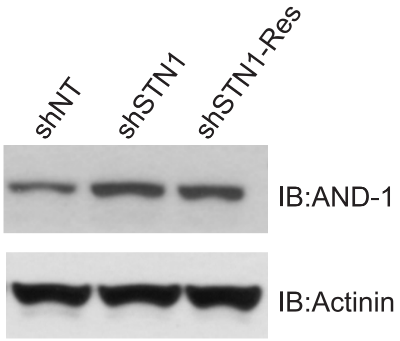


**Figure EV5.** Western blot of AND-1 levels in whole cells lysates in HeLa cells. Actinin was used as a loading control.
